# Ultra-efficient High Resolution 3D Reconstruction of Spatial Omics Data with Neural Transcriptomic Field

**DOI:** 10.64898/2026.05.28.726140

**Authors:** Yuqiao Gong, Xin Yuan, Ruitian Gao, Jinmiao Chen, Zhangsheng Yu

## Abstract

Biological tissues are inherently three-dimensional (3D) ecosystems where spatial architecture dictates cellular function. While spatial omics technologies have revolutionized molecular profiling, they are largely restricted to isolated two-dimensional (2D) tissue sections. Existing computational methods attempting to reconstruct 3D volumes from sparse slices rely heavily on local slice-to-slice interpolation, struggling to balance high-fidelity reconstruction, noise reduction, and atlas-scale efficiency. Here, we present Neural Transcriptomic Field (NTF), a deep learning framework employing multi-resolution hash-grid encoding and implicit neural representations. Unlike interpolation-based approaches that merely bridge adjacent observations, NTF learns a global, continuous 3D representation of the tissue. By modeling the underlying latent biological patterns, NTF intrinsically decouples true molecular signals from technical artifacts, naturally enabling robust denoising and high-fidelity reconstructions. This global field paradigm shatters traditional scalability limits: NTF achieves up to a 1,000× speedup over existing methods, notably reconstructing a 100-million-cell scale 3D whole-mouse embryo atlas in under 15 minutes. Furthermore, NTF can generate super-resolved volumes from sparse input (e.g., utilizing only 10% of slices) and robustly extrapolating into unseen tissue regions. We demonstrate NTF’s versatility across diverse transcriptomic and proteomic datasets, capturing complex spatiotemporal dynamics in *Drosophila* and mouse embryogenesis, and mapping intra-tumoral functional gradients in human breast cancer. Ultimately, NTF provides an unprecedentedly fast, scalable, and robust computational engine for constructing the next generation of comprehensive 3D tissue atlases.

## Introduction

Spatial omics technologies have transformed the study of tissue architecture by measuring molecular states while preserving spatial context ^1-5^. From the original spatial transcriptomics array design to modern sequencing-based and imaging-based platforms, the field has rapidly expanded in resolution, throughput and modality ^2-6^. This growth has created a pressing demand for computational methods that can move beyond descriptive two-dimensional (2D) analysis and recover spatial organization in three dimensions, where developmental gradients, laminar structures, invasive tumor fronts and long-range tissue interactions are often most apparent.

True three-dimensional (3D) measurements are increasingly feasible, but they are still not routine across tissues, modalities and scales. Experimental advances such as STARmap, Open-ST and Deep-STARmap show that direct 3D profiling can reveal spatial structures that are inaccessible in isolated 2D sections ^7-9^. Even so, thick-tissue measurements remain technically demanding, technology-specific and often constrained by panel size, throughput, tissue handling or cost ^3-5,7-9^. As a result, most practical studies still rely on serial or sparsely sampled planar sections and require computational reconstruction to recover missing spatial context.

A broad ecosystem of computational methods has emerged for this purpose ^10^. One class of methods focuses on alignment and integration of adjacent sections, including PASTE, PASTE2, STalign, GPSA and STAligner ^11-15^. A second class aims at explicit 3D reconstruction or atlas building from serial slices, exemplified by SpatialZ and the recent FEAST framework ^16,17^. These methods have substantially advanced the field, but they also expose a persistent trade-off. Alignment-centered approaches are effective for establishing correspondence across slices, yet they do not by themselves provide a continuous high-resolution representation. Reconstruction methods based on virtual-slice synthesis can bridge missing planes, but they are typically driven by local interpolation between observed sections, may inherit observed noise, and can become computationally heavy when applied to atlas-scale data ^16,17^. With the advent of high-resolution platforms like Xenium and Visium HD, a single 2D slice routinely captures hundreds of thousands of spatial elements ^18-20^. Consequently, even a sparse set of input sections easily aggregates to millions of data points, and their subsequent high-resolution 3D reconstructions naturally scale to tens of millions. As spatial omics technologies continue to evolve and the demand for fine-grained resolution grows, high computational efficiency has become an absolute necessity for the practical utility of any 3D reconstruction algorithm.

To overcome the fundamental limitations of local slice-to-slice interpolation and meet the pressing demand for atlas-scale efficiency, we present Neural Transcriptomic Field (NTF). NTF is a unified deep learning framework to reconstruct high-fidelity 3D spatial omics volumes from sparse 2D sections. Rather than relying on discrete interpolation, NTF addresses the unique computational and biological complexities of spatial omics through a series of targeted, parallel architectural innovations: First, to break free from the constraints of discrete local slice interpolation and enable continuous volume generation, we conceptualize the tissue as a global molecular field. Inspired by implicit representations in computer vision (e.g., Neural Radius Filed, NeRF) ^21-23^, NTF learns a continuous coordinate-to-feature mapping, providing a biologically meaningful mathematical representation of the entire tissue architecture. Second, to address the massive computational bottleneck of atlas-scale data, NTF employs a multiresolution hash-encoded architecture. This highly optimized data structure bypasses the heavy overhead of traditional volumetric rendering, allowing the model to process tens of millions of spatial coordinates with unprecedented speed. Third, to systematically decouple true biological signals from the inherited measurement artifacts prevalent in existing methods, NTF introduces a suite of specialized biological modules. It utilizes learnable section-specific embeddings to dynamically align independent spatial planes and correct batch effects; it explicitly models zero-inflation to address transcriptomic dropouts; and it captures cell-level variation to absorb heteroscedastic technical noise. Finally, to accurately reflect the physical limits of spatial assays, we integrate a Point Spread Function (PSF)-aware spatial sampling strategy that models finite acquisition blur.

By capturing the underlying biological reality rather than raw observational noise, we show that NTF combines high reconstruction fidelity with unprecedented computational efficiency. Not only does it achieve up to a 1,000× speedup compared to existing methods, but it also maintains a remarkably lightweight hardware footprint, consuming less than 10 GB of peak memory on a standard NVIDIA A100 (40 GB) GPU. Fundamentally diverging from traditional methods that rely on local 2D slice-to-slice interpolation, NTF learns a holistic, continuous 3D representation of the entire tissue volume. This global perspective ensures that NTF remains highly robust even when only a small fraction of input sections is available. Because the model captures universal spatial trends rather than merely bridging adjacent planes, it successfully super-resolves low-resolution inputs—a task inherently beyond the capability of interpolation-based baselines—and supports challenging extrapolative predictions into unseen tissue regions.

To demonstrate the transformative potential of our model, we deploy NTF across a diverse array of biological systems. Beyond rigorous validation on simulated and real data benchmarks, we show that NTF accurately recovers intricate spatiotemporal developmental dynamics— such as the migration of transcription factors during *Drosophila* embryogenesis. In clinical spatial proteomics, NTF resolves highly irregular contiguous architectures and uncovers distinct intra-tumoral functional gradients (e.g., proliferation and invasion niches) within human breast cancer. Most notably, we demonstrate its unprecedented scalability by reconstructing a 100-million-cell scale, 3D whole-mouse embryo atlas in under 15 minutes, unveiling continuous developmental gradients and the asynchronous nature of mammalian organogenesis. Ultimately, by enabling lightweight in silico sectioning at any desired angle and resolution directly from the trained model, NTF provides a highly versatile and robust computational engine for the next generation of 3D spatial biology.

## Results

### Overview of the NTF model

NTF conceptualizes spatial omics measurements as discrete samples drawn from an underlying continuous 3D molecular field (Fig. 1a, Methods). Within this framework, each spatial observation is defined by its 3D coordinate, its measured molecular profile, and its section identity. To process this data, spatial coordinates are first passed through a multiresolution hash-grid encoding, which is subsequently decoded by a lightweight multi-layer perceptron (MLP) to predict expression values at arbitrary spatial locations. In fundamental contrast to virtual-slice generation pipelines that explicitly synthesize discrete intermediate sections, NTF mathematically learns the holistic field itself. Consequently, once trained, this single, compact representation can be queried at any point in continuous space and rendered at any desired output resolution.

**Fig. 1:**
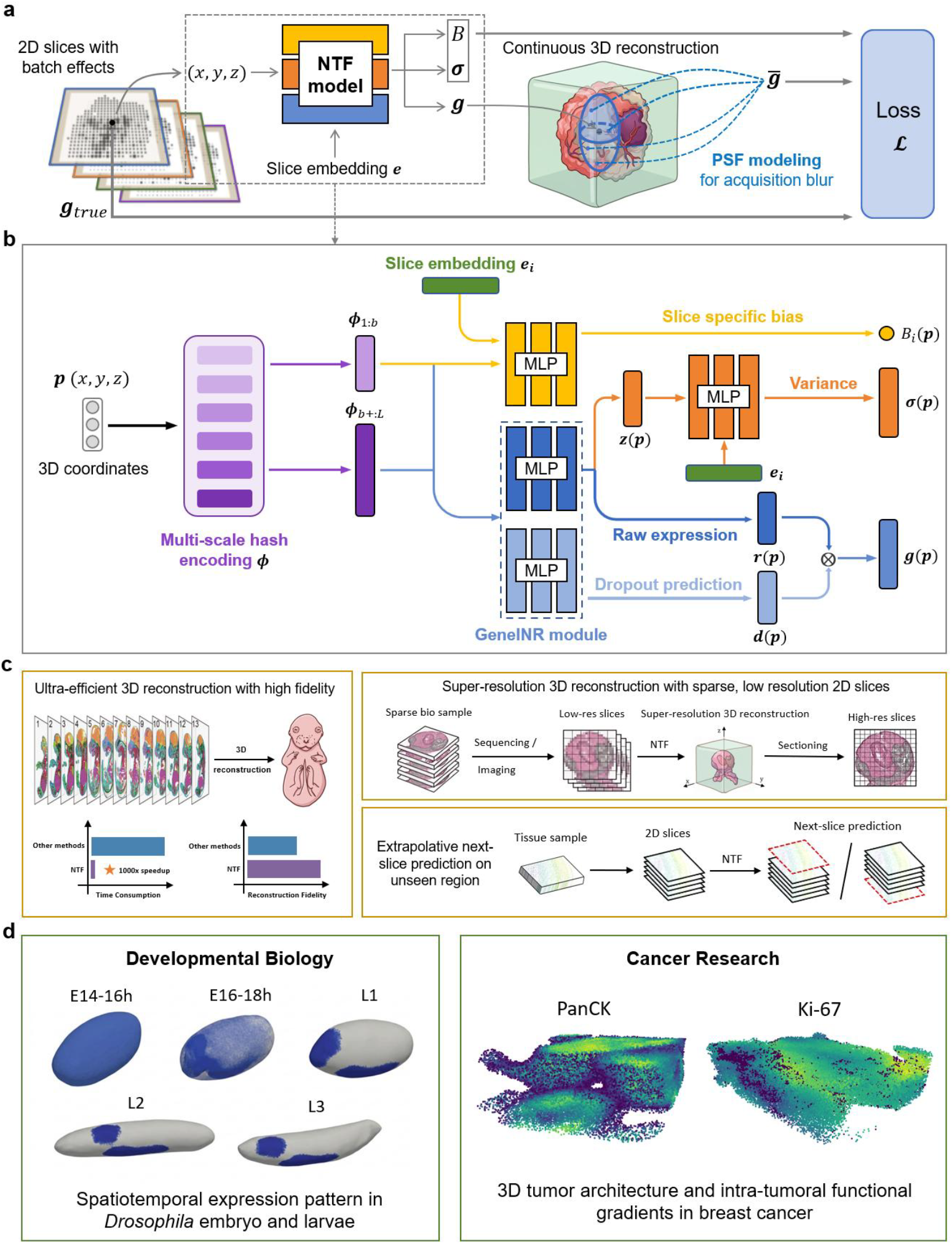
Overview, capabilities, and downstream applications of Neural Transcriptomic Field. **a**, NTF reconstructs a continuous 3D molecular field directly from sparse, 2D spatial omics sections. During training, the model accounts for the finite physical resolution of spatial technologies by aggregating local neighborhood predictions (PSF modeling) to match the observed 2D measurements. **b**, Detailed architecture of the NTF model. Input 3D coordinates are first processed through a multi-scale hash encoding to generate spatial embeddings, which are partitioned into low- and high-frequency components. The low-frequency embedding, combined with a slice-specific embedding (*e*_*i*_), is used to estimate section-dependent bias (to correct for batch effects). The complete embedding drives the GeneINR module to predict raw gene expression (*r*) and calculate a dropout probability (*d*) to address zero-inflation, generating the final expression *g*. Meanwhile, a latent representation (*z*) and the slice embedding jointly model the spatial variance (σ), ensuring robust learning against section-specific technical noise. **c**, Highlighted computational advantages of NTF: achieving ultra-efficient, high-fidelity 3D reconstruction; enabling super-resolution volumetric generation from sparse, low-resolution 2D inputs; and supporting robust extrapolative prediction into unseen spatial regions. **d**, Showcase of biological applications across diverse spatial omics modalities. NTF effectively elucidates complex spatiotemporal expression dynamics during Drosophila embryonic and larval development (developmental biology), and resolves irregular 3D tumor architectures alongside intra-tumoral functional gradients in human breast cancer (spatial proteomics and tumor research).

Beyond this core continuous representation, NTF is natively engineered to decouple true biological signals from the profound technical confounders inherent in spatial assays (Fig. 1b). First, to systematically resolve the prevalent issue of zero-inflation in spatial transcriptomics, NTF features a dedicated dropout module that explicitly predicts the probability of a gene being undetected. Second, to account for uneven capture efficiency and technical noise across serial tissue sections, the architecture introduces learnable slice-specific variables: a bias component to correct for batch effects (section-dependent intensity shifts), and a variance module to absorb cell-level heteroscedastic measurement noise. Finally, to accurately reflect the physical limitations of spatial capture technologies (e.g., spots spanning multiple cells), NTF explicitly models the Point Spread Function (PSF). Rather than treating each coordinate as a dimensionless perfect point, the model aggregates local 3D neighborhood signals to approximate the true physical capture volume before computing the loss against the observed 2D measurements (Fig. 1a).

During training, NTF optimizes a composite objective that explicitly separates gene detection (handling dropouts) from expression quantification. Specifically, the model learns to classify zero versus non-zero observations using a binary cross-entropy loss, while the actual non-zero expression values are fitted using a statistical approach that accounts for varying levels of measurement noise across the tissue (heteroscedastic Gaussian loss). Because biological tissues naturally exhibit spatial continuity, we also incorporate a smoothness regularizer. This constraint encourages neighboring coordinates to share similar expression profiles, effectively suppressing random technical noise while preserving genuine anatomical structures. Notably, despite the massive scale of high-resolution 3D reconstruction, NTF achieves exceptionally fast training and inference speeds. By leveraging the continuous neural field framework augmented with multi-resolution hash encoding and InstantNGP optimizations by NVIDIA ^23^, the model can efficiently process millions of spatial coordinates with minimal computational overhead.

Empowered by these architectural and optimization advances, NTF delivers a suite of powerful computational capabilities (Fig. 1c). Beyond achieving ultra-efficient, high-fidelity 3D reconstruction, the model uniquely enables super-resolution volumetric generation from highly sparse or low-resolution 2D inputs. Furthermore, traditional interpolation methods are mathematically restricted to estimating values between local adjacent observed planes. In fundamental contrast, NTF learns a holistic 3D representation of the entire tissue volume. By capturing global biological trends rather than merely bridging local gaps, this unified continuous field uniquely empowers NTF to perform robust extrapolative predictions into entirely unseen tissue regions extending beyond the observed boundaries. Crucially, these computational strengths translate directly into profound biological insights across diverse spatial omics modalities (Fig. 1d). In the realm of developmental biology, NTF successfully captures intricate spatiotemporal expression dynamics during *Drosophila* embryogenesis and larval development. In clinical tumor research, applying NTF to spatial proteomics data resolves highly irregular 3D tumor architectures and uncovers distinct intra-tumoral functional gradients within human breast cancer. Together, these showcases establish NTF as a highly versatile and scalable engine for decoding complex 3D biological systems.

### NTF achieves reconstruction fidelity with atlas-scale computational efficiency

We first evaluated NTF on simulated data across increasing spatial scales, comparing its performance against SpatialZ and FEAST, two recent methods designed for 3D reconstruction from planar spatial data ^16,17^.. Across all tested scales, NTF consistently achieved the lowest root-mean-square error (RMSE) alongside the highest 3D structural similarity index (SSIM) and Spearman correlation (Fig. 2a, Supplementary Table 1). While SpatialZ served as the strongest numerical baseline, it remained slightly inferior to NTF, whereas FEAST exhibited lower overall reconstruction accuracy. These findings suggest that learning a continuous coordinate-based field is more effective at recovering the true latent spatial signal than relying on explicit slice interpolation.

**Fig. 2:**
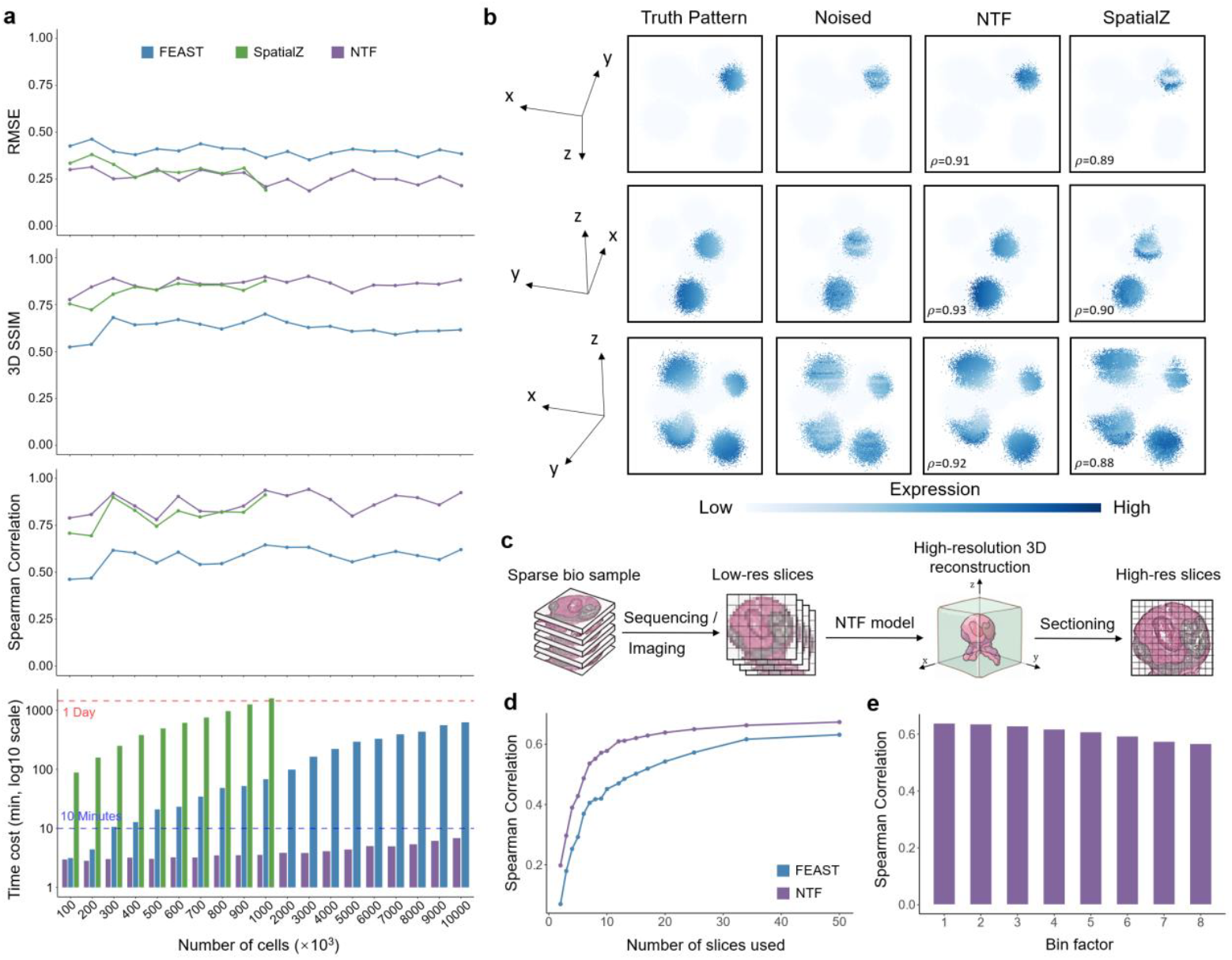
Simulation benchmarks show that NTF combines accuracy, denoising and scalability. **a**, Reconstruction accuracy and runtime across simulated datasets of increasing scale. Full results can be seen in Supplementary Table 1. All methods were benchmarked on a single 40 GB A100 GPU. **b**, Qualitative 3D visualization comparing the reconstructed spatial patterns of three representative genes at the 1-million-cell scale. **c**, Schematic illustrating the sparse-slice and super-resolution reconstruction experimental settings. **d**, Reconstruction performance as a function of the number of provided input slices, evaluated on a 100-slice simulated mouse brain volume. Full results can be seen in Supplementary Table 2. **e**, Performance evaluation when 20 input slices are progressively downsampled in-plane prior to reconstructing the original high-resolution 3D volume. A bin factor of *n* indicates that an *n* × *n* grid of original cells is merged into a single large bin; thus, a larger bin factor corresponds to a lower input resolution.

We next assessed computational scalability. When benchmarked on a single 40 GB A100 GPU, SpatialZ required over 24 hours to process a dataset of 1 million cells, prohibiting its application to larger scales. FEAST reduced runtime relative to SpatialZ but did not match the reconstruction accuracy of NTF. In stark contrast, NTF successfully reconstructed a 10-million-cell dataset in under 7 minutes (Fig. 2a). These results demonstrate that NTF can maintain reconstruction accuracy while scaling to atlas-scale spatial datasets with practical runtimes.

Given that the numerical performance gap between NTF and SpatialZ was modest in certain settings, we next evaluated their qualitative reconstructions for three representative genes using the 1-million-cell simulation benchmark. Because these synthetic observations were generated by adding noise to a known ground-truth pattern, they provided an ideal test for denoising capabilities. NTF recovered spatial expression patterns that closely matched the ground truth rather than the noisy observations, highlighting the robust denoising effects of continuous-field learning combined with smoothness regularization (Fig. 2b). Conversely, SpatialZ tended to reproduce the noisy input structures, indicating that direct slice interpolation fails to recover the underlying, noise-free biological signal.

Finally, we investigated the minimal input information required for accurate high-resolution 3D reconstruction. Using a simulated 2-million-cell mouse brain data originally comprising 100 slices, we systematically subsampled the number of input sections. NTF achieved optimal and stable performance using as few as 10 input slices. In contrast, FEAST required nearly 40 slices to approach its peak accuracy, which still fell short of NTF’s performance (Fig. 2d, Supplementary Table 2). We further evaluated super-resolution reconstruction by starting from 20 slices and progressively reducing their in-plane resolution before reconstructing the original high-resolution 3D volume. A bin factor of n indicates that an *n* × *n* grid of original cells is merged into a single large bin, thus a larger bin factor corresponds to a lower input resolution. NTF showed little degradation even at a bin factor of 8, indicating that the learned field can recover fine-scale structure from coarse inputs when the global spatial organization is retained (Fig. 2e). Neither SpatialZ nor FEAST natively addressed this high-resolution-from-low-resolution setting. Therefore, this analysis was used to assess an additional capability of NTF rather than as a direct like-for-like comparison. Additional qualitative comparisons in Extended Data Fig. 1 confirmed that NTF preserved both high resolution 2D and 3D patterns under sparse and low-resolution input regimes.

### NTF reconstructs missing and boundary slices in experimentally measured 3D mouse visual cortex

We next evaluated NTF on a real 3D spatial transcriptomics dataset from mouse visual cortex generated by STARmap, which provides an experimentally measured volumetric ground truth for systematic validation ^7,16^. We divided the 3D tissue block into seven contiguous slices and designed three reconstruction tasks of increasing difficulty: intermediate-slice prediction, large-block prediction, and boundary-slice prediction.

In the first setting, we trained on slices 1, 3, 5 and 7 and reconstructed the held-out intermediate slices 2, 4 and 6. NTF outperformed both SpatialZ and FEAST when metrics were aggregated across reconstructed genes, achieving a mean Spearman correlation of 0.48, which represents a >10% improvement over competing methods (Fig. 3b). To examine whether these quantitative gains corresponded to improved recovery of biologically structured patterns, we visualized *Rorb*, a laminar marker with a clear layer-restricted expression pattern in cortex ^24^. NTF recovered a sharper and more coherent laminar structure than either baseline, suggesting that NTF preserves layer-restricted spatial structure in held-out sections rather than simply producing a blurred average of neighboring planes. (Fig. 3c). SpatialZ preserved a broader expression pattern with less defined boundaries, whereas FEAST reconstructed more diffuse and less spatially coherent patterns. We then increased the missing gap by training only on the outer slices 1, 2, 6 and 7 while holding out the larger contiguous block of slices 3, 4 and 5. Even under this wider-gap reconstruction regime, NTF remained consistently superior in aggregate metrics and qualitative pattern recovery (Fig. 3e, f). Visualization of other genes shows similar results (Supplementary Figs. 1-2).

**Fig. 3:**
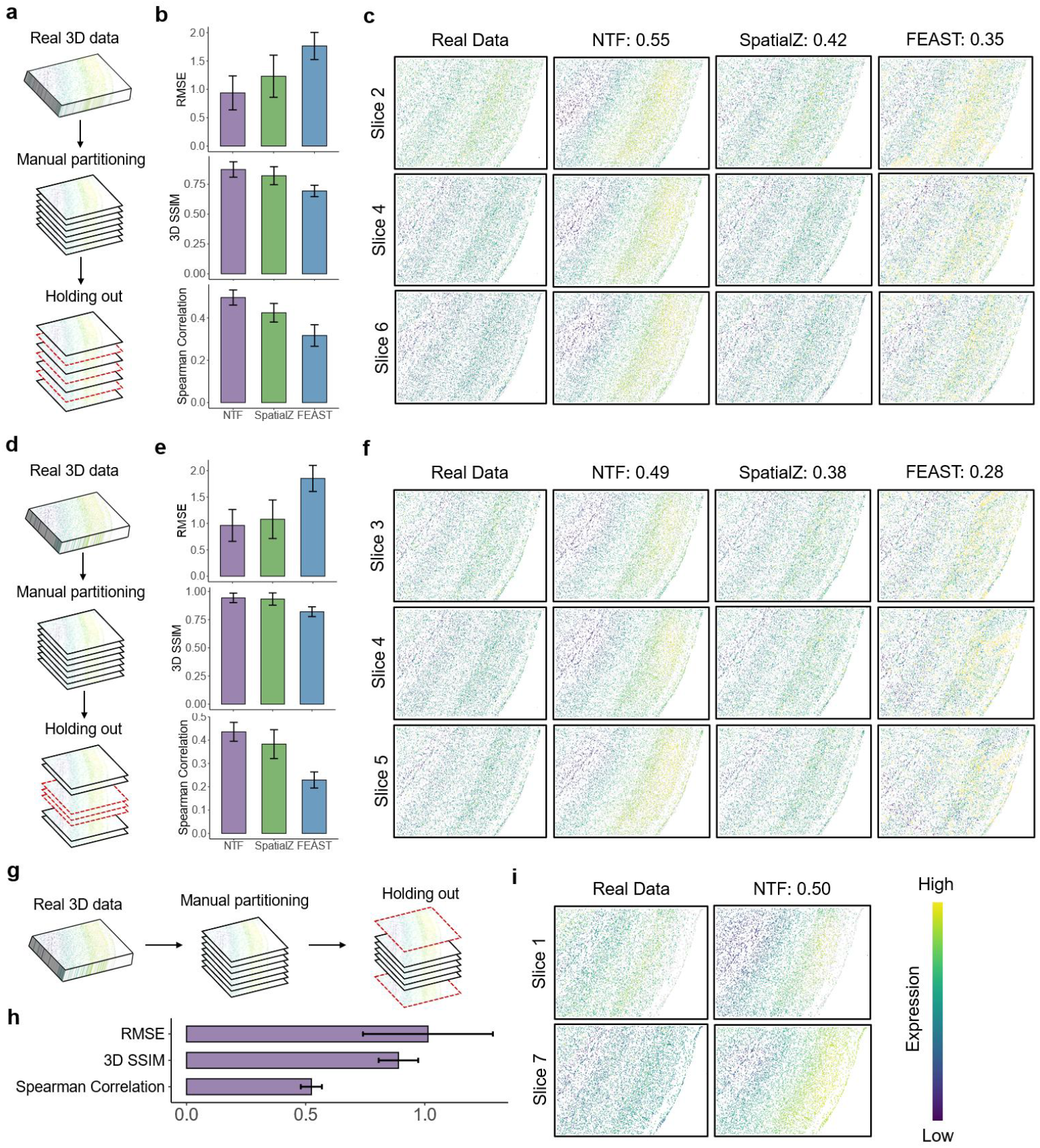
NTF reconstructs missing and extrapolated sections in real 3D mouse visual cortex. **a**, In the first setting, we trained on slices 1, 3, 5 and 7 and reconstructed the held-out intermediate slices 2, 4 and 6 (held-out slices are marked as red dashed line, which is consistent across other scenarios). **b**, Aggregate reconstruction metrics for the first setting. Error bar indicates the standard deviation across all genes. Full results can be seen in Supplementary Table 3. **c**, Representative reconstruction of the laminar marker *Rorb*, showing sharper and more coherent layer-specific recovery by NTF than by SpatialZ or FEAST. **d**, Wider-gap held-out strategy in which slices 1, 2, 6 and 7 are used for training and slices 3, 4 and 5 are reconstructed. **e-f**, Quantitative and qualitative results for the wider-gap setting. **g**, Extrapolative next-slice prediction setting in which slices 2-6 are used for training and slices 1 and 7 are predicted. **h-i**, Reconstruction performance and representative qualitative example of NTF for next-slice prediction.

Because NTF learns a holistic 3D representation of the entire tissue volume rather than merely interpolating between local adjacent planes, it uniquely possesses the capability to project spatial patterns beyond the observed boundaries. Therefore, we also tested extrapolative next-slice prediction by training on the central slices 2–6 and reconstructing the two outermost slices 1 and 7 with NTF. In practice, SpatialZ and FEAST could not handle this setting, whereas NTF could still infer plausible and quantitatively competitive outer slices (Fig. 3h, i, Supplementary Fig. 3). To push the boundaries of this extrapolative capability, we further challenged NTF with more severe, asymmetrical prediction tasks: forecasting two consecutive unseen slices at either end of the tissue block. Specifically, we trained the model on slices 3–7 to predict slices 1 and 2, and conversely, trained on slices 1–5 to predict slices 6 and 7. NTF maintained robust predictive performance even when extrapolating further away from the observed volume boundary (Extended Data Fig. 2, Supplementary Figs. 4-5).

**Fig. 4:**
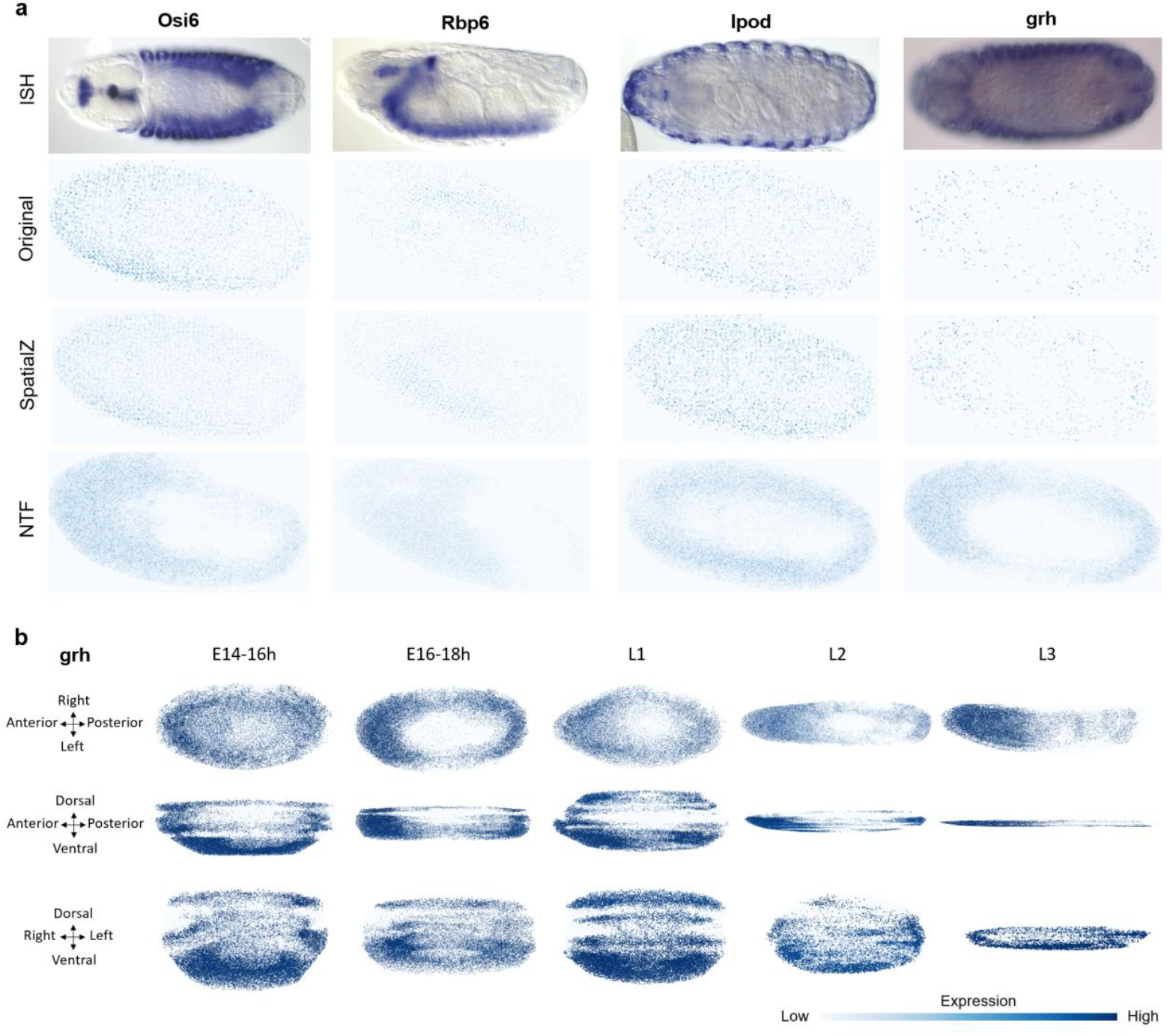
NTF recovers biologically interpretable patterns from low-resolution Drosophila pseudo-3D stacks. **a**, BDGP in situ hybridization references of embryos at stage 13–16 across different genes and 2D projection from original pseudo-3D and reconstructed 3D data by different methods in E14–16 or E16–18 samples. **b**, NTF 3D reconstructions of grh across developmental stages in different 3D views.

Together, these experiments indicate that NTF not only achieves higher reconstruction accuracy than SpatialZ and FEAST in intermediate- and lagre-block held-out slice prediction, but also enables boundary-slice prediction through its learned continuous 3D spatial expression field. This allows NTF to infer unmeasured tissue regions adjacent to the observed volume, a setting that interpolation-based methods cannot address by construction.

### NTF resolves developmental expression programs from low-resolution *Drosophila* pseudo-3D stacks

To assess whether NTF can recover biologically meaningful 3D patterns from low-resolution stacked spatial data, we applied it to a publicly available Drosophila embryo and larval pseudo-3D dataset derived from serial Stereo-seq sections ^25^. We benchmarked the reconstruction fidelity against independent in situ hybridization (ISH) images of stages 13–16 embryos from the Berkeley Drosophila Genome Project (BDGP) ^26^. To ensure cross-modality comparability, 3D reconstructed volumes were projected onto 2D planes along z axis using summed expression as in original research ^25^. Here we projected results from embryonic stages E14-16 or E16-18 onto 2D planes for qualitative comparison. Across four genes with distinct spatial expression patterns (*Osi6, Rbp6, Ipod* and *grh*), the 2D projections of NTF reconstructions reproduced the corresponding ISH patterns, including the peripheral signal of *Osi6, Ipod* and *grh* and the spatial asymmetries, anterior enrichment of *Rbp6*. As FEAST consistently underperformed compared to NTF and SpatialZ in our preceding quantitative benchmarks, we focused this detailed visual evaluation exclusively on SpatialZ. In contrast to NTF, the raw Stereo-seq stack and SpatialZ reconstructions retained the sparsity of the input and did not resolve these spatial features (Fig. 4a). Because SpatialZ relies heavily on local adjacent-slice interpolation and lacks global cross-slice constraints, it is vulnerable to the noise and morphological discrepancies present in the raw sections. Consequently, rather than recovering biological boundaries, the SpatialZ reconstructions remained visually diffuse and constrained by the coarse input.

We next examined whether NTF could recapitulate stage-dependent changes in spatial expression using the transcription factor *grh* (grainy head), which shows a marked and well-characterized shift in localization across embryonic and larval stages ^27,28^. This made *grh* a useful test case for assessing whether NTF can recover developmentally regulated spatial reorganization. In the embryonic stages (E14-16 h and E16-18 h), the NTF-reconstructed *grh* signal showed a shell-like epidermal pattern, consistent with its known association with epidermal barrier and cuticle formation ^27^. In the larval stages (L1–L3), the reconstructed signal showed enriched expression in the brain lobes and ventral nerve cord, consistent with previous reports linking Grainy head to post-embryonic neuroblast regulation and neural proliferation programs ^28^ (Fig. 4b).These stage-dependent changes in spatial organization were less clearly resolved in the original low-resolution stack and were only partially recovered by SpatialZ (Extended Data Fig. 3), suggesting that NTF can recapitulate developmentally regulated spatial patterns that are otherwise obscured by input sparsity.

### NTF reconstructs 3D tumor architecture and intra-tumoral functional gradients from spatial proteomics

To evaluate whether NTF extends beyond transcriptomic data, we applied it to a human breast cancer dataset profiled via imaging mass cytometry (IMC) ^29^. Compared with the transcriptomic datasets analyzed above, this spatial proteomic dataset provides a more heterogeneous and structurally complex test case. Using panCK as an epithelial tumor marker, NTF reconstructed a continuous 3D protein field that produced more coherent tumor boundaries and morphology than the naive pseudo-3D assembly (Fig. 5a-c). Regions that appeared disconnected in individual 2D sections were reconstructed as spatially contiguous extensions of a single irregular epithelial tumor structure. These results suggest that NTF can recover continuous 3D tumor morphology from sparsely sampled spatial proteomics data.

**Fig. 5:**
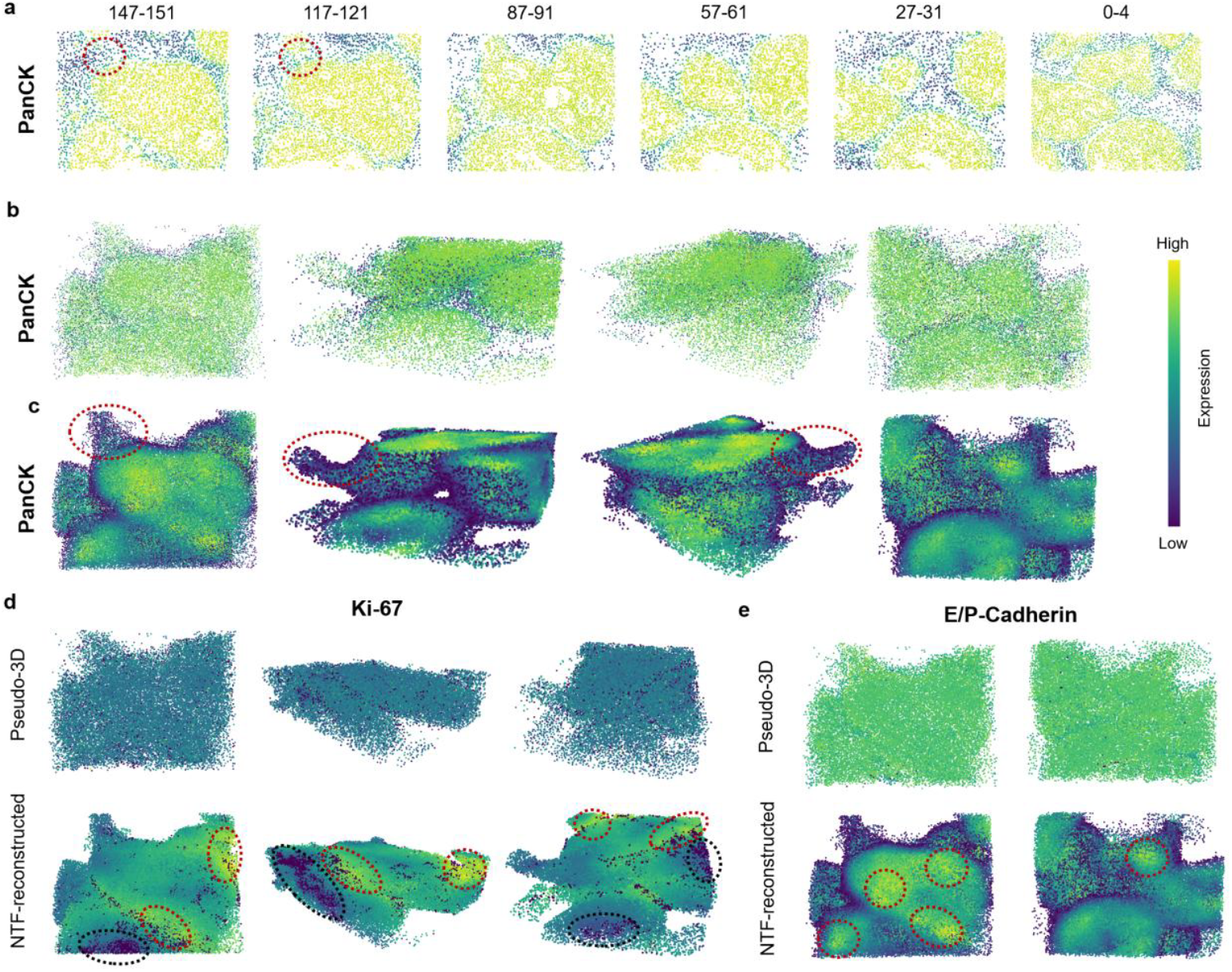
NTF reconstructs complex 3D architecture and functional heterogeneity in human breast cancer from sparse IMC slices. **a**, Representative 2D IMC slices of human breast cancer across depth. Red dashed circle indicates a continuous tumor region which can be seen as disconnected tumors in 2D view. **b**, Different view of pseudo-3D volume formed by direct stacking of the 2D slices. **c**, Different view of NTF reconstruction of the panCK protein expression, resolving tumor morphology, boundaries and a contiguous 3D structure that is difficult to infer from 2D views alone (Red dashed circle). **d**, Comparison of Ki-67 (proliferation marker) expression between the pseudo-3D assembly (top) and the NTF 3D reconstruction (bottom). Red dashed circles indicate highly expressed centers while black dashed circles indicate lowly expressed centers. **e**, 3D spatial distribution of E/P-Cadherin (invasion and migration markers).

We next examined whether NTF could recover spatially structured distributions of functionally relevant protein markers within the tumor volume. We focused on Ki-67, a marker of cellular proliferation, and E/P-Cadherin, which has been associated with tumor invasion and cell migration ^30,31^. In the naive pseudo-3D assembly, discontinuities between adjacent sections made the spatial organization of these markers difficult to interpret (Fig. 5d, e). By contrast, NTF reconstructed spatially coherent 3D protein distributions for both markers. For Ki-67, multi-angle visualization of the reconstructed volume revealed two regions with elevated proliferative signal (red dashed) and two regions with relatively low signal (black dashed, Fig. 5d), suggesting that NTF can identify spatially organized proliferative heterogeneity within the tumor mass. For E/P-Cadherin, NTF revealed a boundary-enriched pattern relative to the tumor core (Fig. 5e), consistent with reported boundary enrichment of Cadherin in invasive breast cancer ^31^.Together, these results suggests that NTF can recover continuous 3D morphology and spatially structured protein expression from noisy, sparsely sampled clinical sections, extending its applicability to spatial proteomics.

### Rapid atlas-scale 3D reconstruction of whole-embryo spatial transcriptomics

To evaluate NTF’s scalability and its ability to capture holistic developmental programs, we applied it to a late-stage (E16.5) mouse embryo dataset from the Mouse Organogenesis Spatiotemporal Transcriptomic Atlas (MOSTA), comprising 13 serial spatial sections^32^. Remarkably, despite simultaneously reconstructing over 1,000 genes, NTF generated a high-resolution, 100-million-voxel scale 3D whole-embryo atlas in less than 15 minutes on a single 40GB A100 GPU, with peak GPU memory usage < 10GB (Fig. 6a). This unprecedented efficiency establishes NTF as a practical tool for atlas-scale spatial omics.

**Fig. 6:**
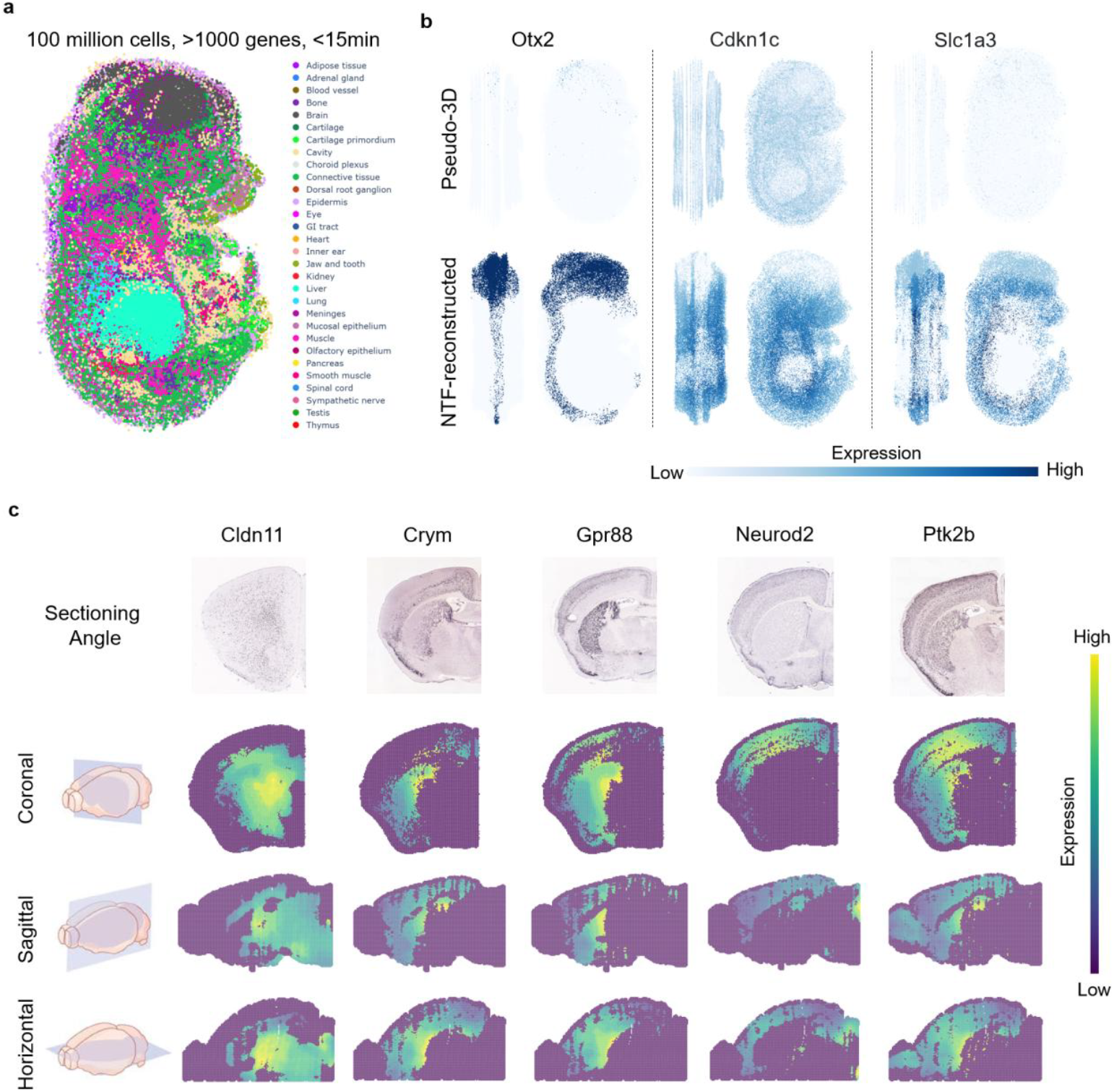
NTF generalizes to large embryonic reconstructions and enables light-weighted in silico sectioning. **a**, High-resolution 3D reconstruction of a mouse embryo cell atlas containing 10 million cells with more than 1,000 genes. **b**, Qualitative comparison of 3 genes between the raw pseudo-3D assembly (top) and the NTF reconstruction (bottom). **c**, Reference in situ hybridization images of coronal sections from the Allen Mouse Brain Atlas and NTF in silico section results from different section angles.

We next compared the spatial expression of developmental regulators between the raw pseudo-3D stack and the NTF reconstruction (Fig. 6b). While the raw stack exhibited slice-to-slice discontinuities and technical noise, NTF recovered continuous organ boundaries and spatial gradients. For instance, the NTF reconstruction clearly delineated the localized expression of *Otx2* in the developing eyes, brain, and spinal cord regions. Furthermore, NTF resolved the spatial heterogeneity of *Cdkn1c* across different tissues. The reconstructed volume demonstrated enriched *Cdkn1c* expression in developing skeletal muscle, contrasting with diminished expression in early-developing organs such as the brain, heart, and liver. This organ-specific divergence was largely obscured by discontinuities in the raw pseudo-3D stack. Finally, NTF recovered continuous expression gradients along anatomical axes. For *Slc1a3*, the 3D reconstruction revealed a progressive expression gradient across the central nervous system, increasing along the anterior-posterior axis toward the spinal cord. This continuous spatial transition was not resolvable in the discrete pseudo-3D input. Together, these results demonstrate that NTF can synthesize sparse 2D sections into a coherent 3D representation, preserving both structural boundaries and functional gradients.

### NTF enables lightweight in silico sectioning across arbitrary spatial planes and resolutions

A practical advantage of NTF is that virtual sectioning does not require precomputing a dense atlas. Atlas-first workflows such as SpatialZ typically generate a dense reconstructed volume and then resample planes from that atlas for downstream visualization or analysis ^16^. In NTF, the trained field itself is the stored representation. After training, we save the model parameters together with geometry metadata, and any virtual plane can be generated later by directly querying the field at the desired angle, position and output resolution (Extended Data Fig. 4). This reduces both storage and preprocessing requirements and makes interactive sectioning feasible.

To validate this capability, we trained NTF on a mouse brain dataset and generated virtual coronal slices that were compared with reference in situ hybridization images from the Allen Mouse Brain Atlas ^33^. The virtual sections reproduced major spatial expression patterns with close visual agreement to the reference data (Fig. 6c). Because the field is continuous, the same trained model could also generate sagittal and horizontal sections without retraining or atlas resampling, exposing complementary two-dimensional views of the same 3D molecular structure (Fig. 6c). These experiments show that NTF turns sparse 2D input sections into a compact 3D representation that remains directly queryable long after model fitting has finished.

## Discussion

We present Neural Transcriptomic Field (NTF), an ultra-efficient coordinate-based neural field model that reconstructs continuous, high-resolution 3D spatial omics landscapes from sparse planar measurements. By integrating multiresolution hash-grid encoding with explicit observation models for zero inflation, heteroscedastic uncertainty, section-specific effects, and acquisition blur, NTF learns a compact function that can be queried at arbitrary coordinates. This formulation yields state-of-the-art reconstruction fidelity while achieving unprecedented computational scalability, reconstructing 100 million spatial cells in under 15 minutes on a single GPU.

Current computational strategies for 3D spatial omics primarily rely on pairwise slice alignment or explicit interpolation between observed sections. While effective for establishing correspondences, these approaches often inherit measurement noise, struggle with large missing gaps, and incur substantial memory and runtime costs when dense atlases must be computed. NTF addresses these bottlenecks by learning a continuous latent field regularized for spatial smoothness and uncertainty-aware expression prediction. The hash-grid encoder captures multi-scale spatial dependencies efficiently, while the composite loss function separates zero-handling from expression reconstruction, promoting denoised signal recovery rather than mere interpolation of noisy observations. This methodological shift explains NTF’s consistent superiority in structural similarity and correlation metrics, its robustness to aggressive slice subsampling, and its ability to recover fine-scale patterns even from low-resolution inputs.

Beyond quantitative benchmarks, NTF demonstrates broad practical utility across diverse biological contexts and measurement modalities. In real 3D mouse cortex data, NTF accurately interpolates missing slices and, notably, extrapolates to unobserved outer sections—a capability largely inaccessible to interpolation-centric methods. Its application to low-resolution *Drosophila* developmental stacks reveals stage-specific expression dynamics that are otherwise obscured, while successful transfer to imaging mass cytometry and large-scale mouse embryo atlases underscores modality-agnostic flexibility. Crucially, the trained field serves as a lightweight, persistent representation that supports interactive, arbitrary-resolution virtual sectioning without retraining or atlas resampling. This workflow drastically reduces storage overhead and preprocessing latency, making high-resolution 3D spatial analysis accessible for iterative exploration and downstream integration.

Recent perspectives emphasize that malignancies act as organized 3D entities whose physical shapes are intrinsically linked to their biological circuitry and developmental modalities ^34^. Viewed through this lens, the highly complex and interconnected 3D architecture uncovered by NTF is not mere morphological noise, but rather reflects an optimal ‘space-filling’ strategy—reminiscent of branching morphogenesis—used by complex or metastatic breast cancers to maximize epithelial surface area and survival signals from the microenvironment. By resolving these intricate 3D geometries from sparse 2D slices, NTF provides a vital computational tool for decoding the structural basis of tumor progression.

Several limitations and exciting avenues for future development warrant consideration. While NTF successfully leverages holistic 3D representations to perform unprecedented extrapolation beyond observed boundaries, the spatial extent of this extrapolation is naturally constrained by the information density of the sampled volume. Projecting complex molecular fields deeply into distant, un-biopsied macroscopic regions remains a fundamental computational and biological challenge. Future developments will focus on pushing these extrapolative boundaries further, potentially by coupling the NTF continuous field with morphological priors from routine non-destructive imaging (e.g., H&E or MRI) or large-scale spatial foundation models. Expanding this predictive envelope holds profound clinical significance. In clinical pathology and surgical oncology, tissue availability is strictly limited, and extensive serial sectioning is often destructive and impractical. If a model could reliably reconstruct the entire 3D tumor microenvironment—and critically, predict deep invasive tumor margins or hidden metastatic niches—from just a few sparse core-needle biopsies, it would be highly transformative. It would preserve scarce clinical material, minimize tissue waste, and provide oncologists with a comprehensive, virtual 3D molecular map to guide precision surgical resections and targeted therapies.

By providing an open, ultra-efficient, and queryable 3D representation, NTF bridges a critical gap between sparse spatial measurements and continuous tissue-scale modeling. As spatial omics continues to scale in dimension and complexity, NTF offer a flexible foundation for next-generation 3D molecular cartography, ultimately accelerating our understanding of tissue organization in development, homeostasis, and disease.

## Methods

### Neural Transcriptomic Field (NTF) Framework

The Neural Transcriptomic Field (NTF) parameterizes the continuous spatial landscape of gene expression as a coordinate-based neural filed. Let *x* ∈ *R*^3^ denote the continuous 3D spatial coordinates within a tissue. To capture high-frequency spatial variations effectively while maintaining computational efficiency, we map the input coordinates into a high-dimensional feature space using a multi-resolution hash grid encoding γ(*x*).

The core of NTF consists of decoupled Multi-Layer Perceptrons (MLPs) that predict the expected gene expression, observation variance, and dropout probability. Specifically, the GeneINR module *f*(θ) takes the hash-encoded features to predict the latent gene expression *e* ∈ *R*^*G*^ for *G* genes and an intermediate feature vector *z*: (*e, z*) = *f*_θ_(γ(*x*)). To ensure biological validity (non-negativity), the raw expression outputs are passed through a Softplus activation, *µ* = *ln*(1 + *exp*(*e*)).

To account for the finite resolution and optical/physical dispersion inherent in spatial transcriptomics technologies, NTF models the observed signal as a spatial integration over a physical volume defined by a Point Spread Function (PSF). We approximate this integration via Monte Carlo sampling. For a given spatial coordinate *x*, we draw N sub-samples perturbed by Gaussian noise parameterized by the anisotropic resolution in three-dimensional coordinates Σ_**psf**_: *x*^(*i*)^ *∼* 𝒩(*x*, Σ_psf_), *i* ∈ 1, …, *N*. The final predicted expression vector 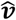 is obtained by marginalizing over these samples:

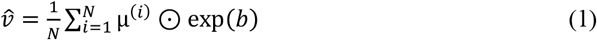

where b is slice- or level-specific bias term predicted by an auxiliary network to correct for batch effects across multiple tissue slices.

### Probabilistic Modeling and Zero-Inflation

Spatial omics data are frequently characterized by high sparsity and zero-inflation. To explicitly model the technical dropouts (where transcripts are present but fail to be captured), NTF employs a parallel dropout network *h*_*ϕ*_ that estimates the logit probabilities of a zero-count observation: *d*^(*i*)^ = *h*_*ϕ*_ (γ(*x*^(*i*)^)) The aggregated dropout logit 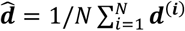 is used to predict the probability of observing a zero strictly due to technical artifacts.

Furthermore, we model the heteroscedastic noise (variance) of the transcriptomic measurements. A variance network *g*_*Ψ*_ is conditioned on the intermediate feature *z* and the slice embedding *s*_*k*_ (for the k-th slice) to output the gene-specific log-variance:

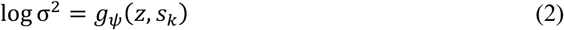

The total predicted variance includes both the pixel-level variance *σ*^**2**^ and a learnable slice-level variance offset.

### Loss Formulation

The NTF model is trained end-to-end by minimizing a composite objective function that handles sparsity, observation uncertainty, and spatial continuity:

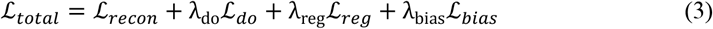

### Uncertainty-Aware Reconstruction Loss (ℒ_recon_)

To robustly fit the expression values without being biased by noise, we formulate the reconstruction loss as the negative log-likelihood of a Gaussian distribution. Notably, to decouple true expression gradients from technical dropouts, this loss is strictly evaluated on the non-zero observations:

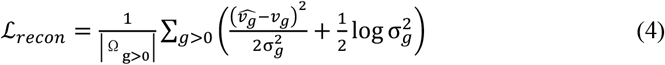

where *v*_*g*_ is the ground-truth expression, 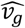 is the predicted expression, and 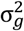 is the predicted variance for gene *g*.

### Zero-Inflation / Dropout Loss (ℒ_do_)

The dropout network is supervised via a weighted Binary Cross-Entropy (BCE) loss against a binary mask y indicating whether the ground-truth expression is zero:

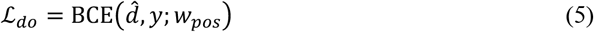

where *w*_*pos*_ represents the ratio of non-zero to zero values, mitigating the extreme class imbalance prevalent in spatial datasets.

### Spatial Smoothness Regularization (ℒ_reg_)

To enforce biologically plausible spatial continuity and prevent overfitting to high-frequency artifacts, we introduce a local smoothness prior. For a subset of sampled points ***x***^(***i***)^, we generate neighboring points uniformly distributed within a local radius *r*(*δ ∼ U*(−*r, r*)). We penalize the Mean Squared Error (MSE) between their predicted expressions:

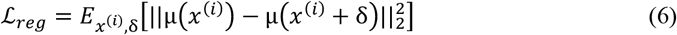

### Bias Regularization (ℒ_bias_)

When slice-level bias modeling is enabled, an L2 penalty 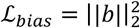 is applied to the log-bias terms to prevent the network from incorrectly attributing true spatial variations to artifactual batch effects.

### Training and Inference

We train NTF with Adam, mixed precision by default, multi-step learning-rate decay and early stopping. The default training configuration uses 20,000 optimization iterations, a batch size of 8,192, a learning rate of 1e-4 and four local coordinate samples per observation. Because training samples observed points directly rather than rasterizing the entire volume, the computational cost scales favorably to large datasets. At inference time, the trained field is queried directly at new coordinates to produce reconstructed expression.

### Evaluation Metrics

To comprehensively evaluate the performance of different methods in reconstructing and predicting spatial gene expression, we quantified the accuracy, monotonic concordance, and spatial structural preservation using Root Mean Square Error (RMSE), Spearman correlation, and 3D Structural Similarity Index Measure (3D SSIM), respectively.

Let *N* denote the total number of evaluated spatial coordinates (or cells/voxels). For a given gene, let *v*_*i*_ and 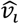 represent the ground-truth and the predicted expression values at the *i*-th spatial location, respectively.

### Root Mean Square Error (RMSE)

RMSE measures the absolute magnitude of the prediction error, penalizing larger deviations. It is defined as the square root of the average squared differences between the predicted and observed expression values:

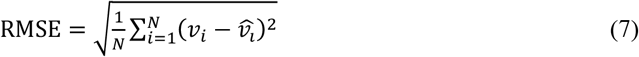

### Spearman Correlation (*r*_*s*_)

Given the inherent noise and nonlinear scaling often present in spatial omics measurements, rank-based correlation is more robust than linear correlation. Spearman’s rank correlation coefficient assesses the monotonic relationship between the true and predicted spatial expression landscapes. Let *R*(*v*_*i*_) and 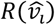 denote the fractional ranks of the raw values *v*_*i*_ and 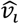among all *N* locations. The Spearman correlation is equivalent to the Pearson correlation of these rank values:

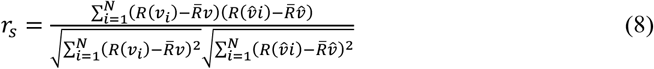

where 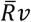 and 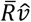 are the mean ranks.

### 3D Structural Similarity Index Measure (3D SSIM)

To evaluate the perceptual quality and the structural fidelity of the predicted 3D continuous gene expression fields, we extended the standard 2D SSIM to 3D volumes. 3D SSIM captures the similarity of spatial expression patterns by jointly assessing luminance (mean expression), contrast (variance), and structure (covariance) within local 3D sliding windows. For two local 3D volumetric windows *x* and *y* (extracted from the ground-truth and predicted 3D expression volumes, respectively), the local SSIM is calculated as:

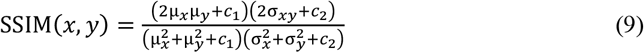

where μ_*x*_ and μ_*y*_ are the local weighted means, 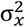 and 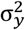 are the local variances, and σ_*xy*_ is the local covariance computed using a 3D Gaussian weighting kernel. *c*_1_ = (*k*_1_*L*)^2^ and *c*_2_ = (*k*_2_*L*)^2^ are stabilization constants to avoid numerical instability, with *L* representing the dynamic range of the expression values. The final overall 3D SSIM score is the mean of the local SSIM values calculated across the entire 3D spatial field.

### Simulation study

Simulation experiments were designed to quantify reconstruction accuracy, computational efficiency, robustness to sparse sectioning and robustness to low input resolution. The scale benchmark compared NTF with SpatialZ and FEAST across increasing dataset sizes up to 10 million cells. To evaluate sparse-input robustness, a simulated 2-million-point mouse brain volume was partitioned into 100 slices, from which progressively fewer sections were retained for training. To evaluate super-resolution reconstruction, 20 slices were retained and their in-plane resolution was reduced by spatial binning before reconstructing the original high-resolution 3D volume.

To systematically quantify reconstruction accuracy, computational efficiency, and robustness against both sparse sectioning and low spatial resolution, we designed a series of in silico experiments. The ground-truth continuous spatial transcriptomic fields were mathematically simulated to reflect the complex spatial patterns, noise profiles, and zero-inflation inherent to real spatial omics datasets.

### Generation of simulated spatial transcriptomic fields

To mimic realistic tissue architectures, cells were assigned to distinct spatial domains. We explicitly modeled both domain-specific differential genes and uniformly expressed housekeeping genes. For differential genes, we mathematically defined diverse underlying spatial expression landscapes within active domains, including linear gradients along the x, y, or z axes, as well as radial distributions (e.g., center-high to periphery-low). To capture the overdispersion of sequencing counts, the ground-truth expression matrix was sampled from a Zero-Inflated Negative Binomial distribution, parameterized by the dropout probability, spatial pattern mean and a predefined dispersion factor. To rigorously simulate technical artifacts, we injected slice-specific batch effects (Gaussian offsets) and global random Gaussian noise.

### Scale benchmark

To evaluate computational scalability and efficiency, we generated datasets of increasing sizes, up to 10 million cells, using the aforementioned spatial pattern generator. Cells were randomly distributed into 3D clusters with overlapping spatial gradients. The performance and runtime of NTF were benchmarked against existing methods, including SpatialZ and FEAST, on these large-scale simulated coordinates.

### Robustness to sparse sectioning

To evaluate the interpolative capacity of NTF across large z-axis gaps, a simulated 2-million-point tissue volume (based on a domain-annotated mouse brain model) was partitioned into 100 consecutive slices. We simulated sparse input scenarios by progressively sub-sampling the training data at varying intervals. The model was trained exclusively on the sparse subsets, and its predictive performance was evaluated by querying the continuous neural representation at the spatial coordinates of all original 100 high-resolution slices, comparing the output against the unobserved ground truth.

### Super-resolution spatial reconstruction

To assess the model’s ability to mathematically super-resolve low-resolution spatial inputs (analogous to array-based technologies like 10x Visium), we retained a subset of 20 slices from the high-resolution simulation and applied a spatial degradation algorithm. We performed in-plane spatial binning by partitioning the coordinates of each slice into 2D grids. The grid size was determined by scaling the average intercellular distance by a designated binning factor. Within each spatial bin, single-cell coordinates were aggregated to their geometric centroid, and their gene expression profiles were summed to simulate bulk-like capture spots. NTF was trained solely on this fully degraded, low-resolution 3D dataset. To evaluate super-resolution performance, the trained continuous field was queried at the exact high-resolution single-cell coordinates of the original dataset, allowing us to quantify how accurately NTF reconstructs fine-grained spatial structures from coarse-grained inputs.

## Data availability

All datasets analyzed for this study are publicly available. The 3D visual cortex dataset, generated using STARmap technology, is available through STARmap Resources at https://www.starmapresources.org/data. Drosophila embryo and larval pseudo-3D stacks from Stereo-seq can be download from https://db.cngb.org/stomics/flysta3d. The human breast cancer IMC data are available on Zenodo via https://doi.org/10.5281/zenodo.4752030. The 13-slice E16.5 mouse embryo dataset from MOSTA is available from https://db.cngb.org/stomics/mosta. Mouse brain data can be accessed and downloaded from https://alleninstitute.github.io/abc_atlas_access/descriptions/Zhuang-ABCA-2.html. The processed and NTF-reconstructed datasets are publicly available via zenodo at https://doi.org/10.5281/zenodo.19590994. Source data are provided with this paper.

## Code availability

The code used to develop the model, perform the analyses and generate results in this study is publicly available and has been deposited in GitHub at https://github.com/GYQ-form/NTF, under MIT license.

## Acknowledgements

This study was supported by grants from the National Natural Science Foundation of China (Grant No. 12171318 to Z.Y., 325B2022 to Y.G. and 32500566 to X.Y.), the Shanghai Science and Technology Commission (Grant No. 24JS2810200, 24DZ2260100, 23XD1401900, 23DZ2290600 to Z.Y. and 25ZR1402274 to X.Y.), the Medical Engineering Cross Fund of Shanghai Jiao Tong University (Grant No. YG2023ZD21 to Z.Y., and YG2026QNA28 to R. G.), the Shanghai Jiao Tong University Start-up Program for Newly Recruited Young Faculty (Grant No. 25X010506038 to X.Y.) and Yu Lab. The computations in this paper were run on the Siyuan-1 cluster supported by the Center for High-Performance Computing at Shanghai Jiao Tong University.

## Authors’ contributions

Y.G. performed the main research, analyzed data, and wrote the original manuscript. X.Y. and Y.G. investigated and interpretated the analysis outcomes. J.C. and Z.Y. supervised the research. Y.G., X.Y., R.G., J.C and Z.Y. discussed and revised the manuscript.

## Extended Data Figure

**Extended Data Figure 1:**
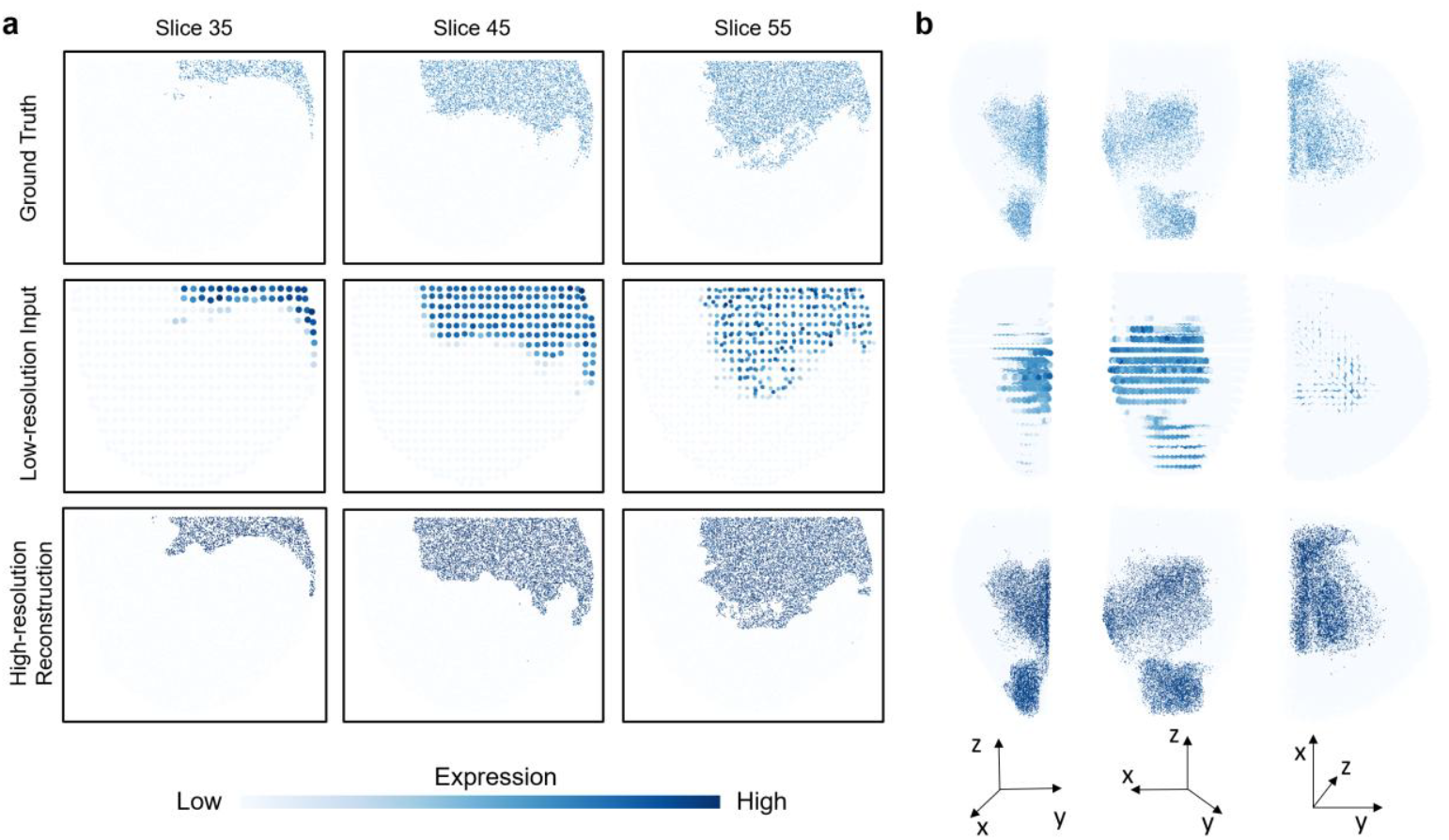
Qualitative robustness of NTF under sparse and low-resolution simulation inputs. **a**, Representative 2D reconstructions from sparse-slice simulations. **b**, Corresponding 3D reconstructions showing preservation of volumetric structure despite aggressive slice subsampling.

**Extended Data Figure 2:**
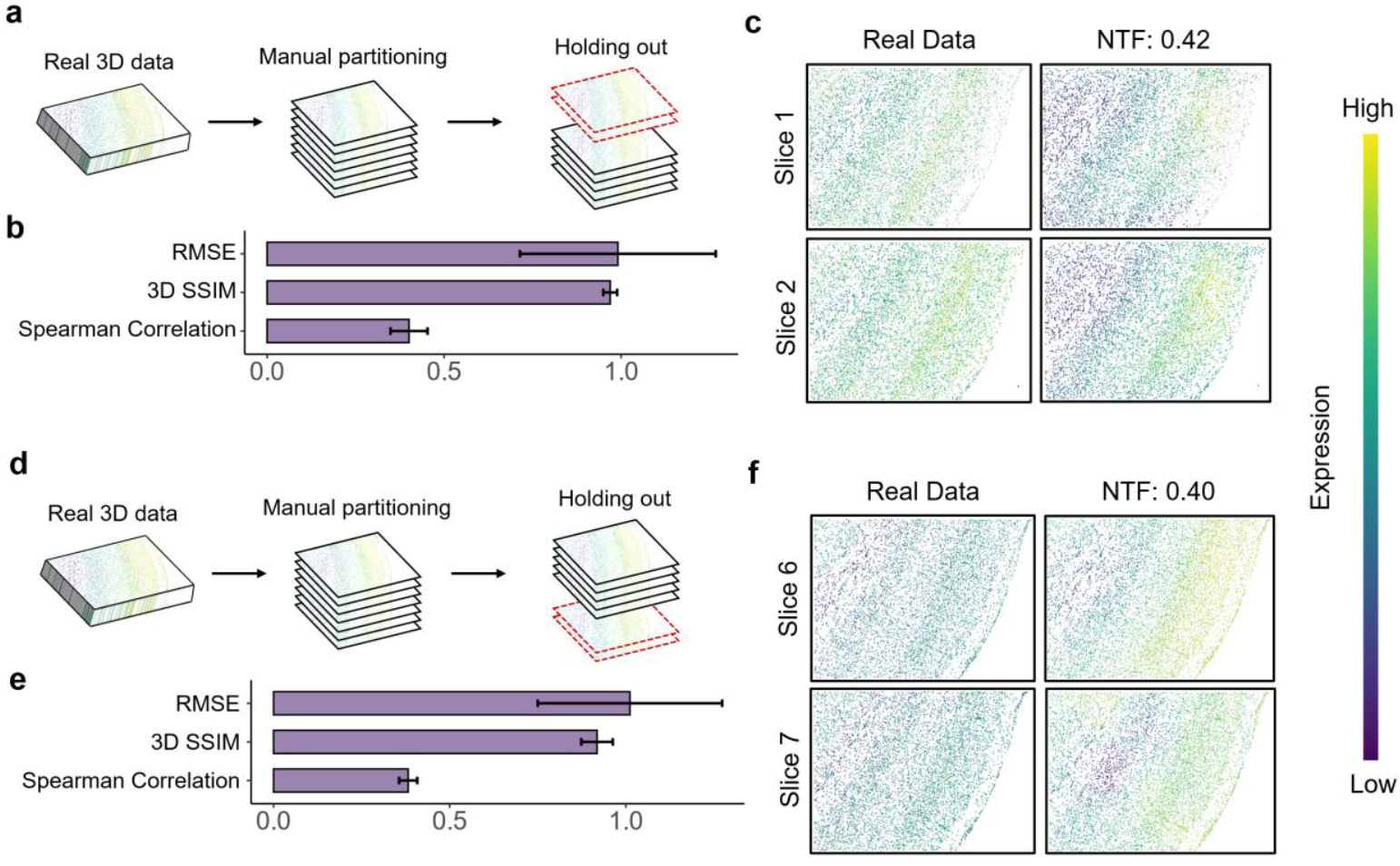
NTF demonstrates robust extrapolative prediction for multiple consecutive unseen slices. **a**, Backward extrapolation: NTF predicts the expression patterns of slices 1 and 2 using only slices 3–7 as training input. **b**, Aggregate reconstruction metrics for all the genes. **c**, Representative reconstruction of the laminar marker Rorb by NTF. **d**, Forward extrapolation: NTF predicts slices 6 and 7 using only slices 1–5 as training input. **e-f**, Reconstruction performance and representative qualitative example of NTF for forward extrapolation.

**Extended Data Figure 3:**
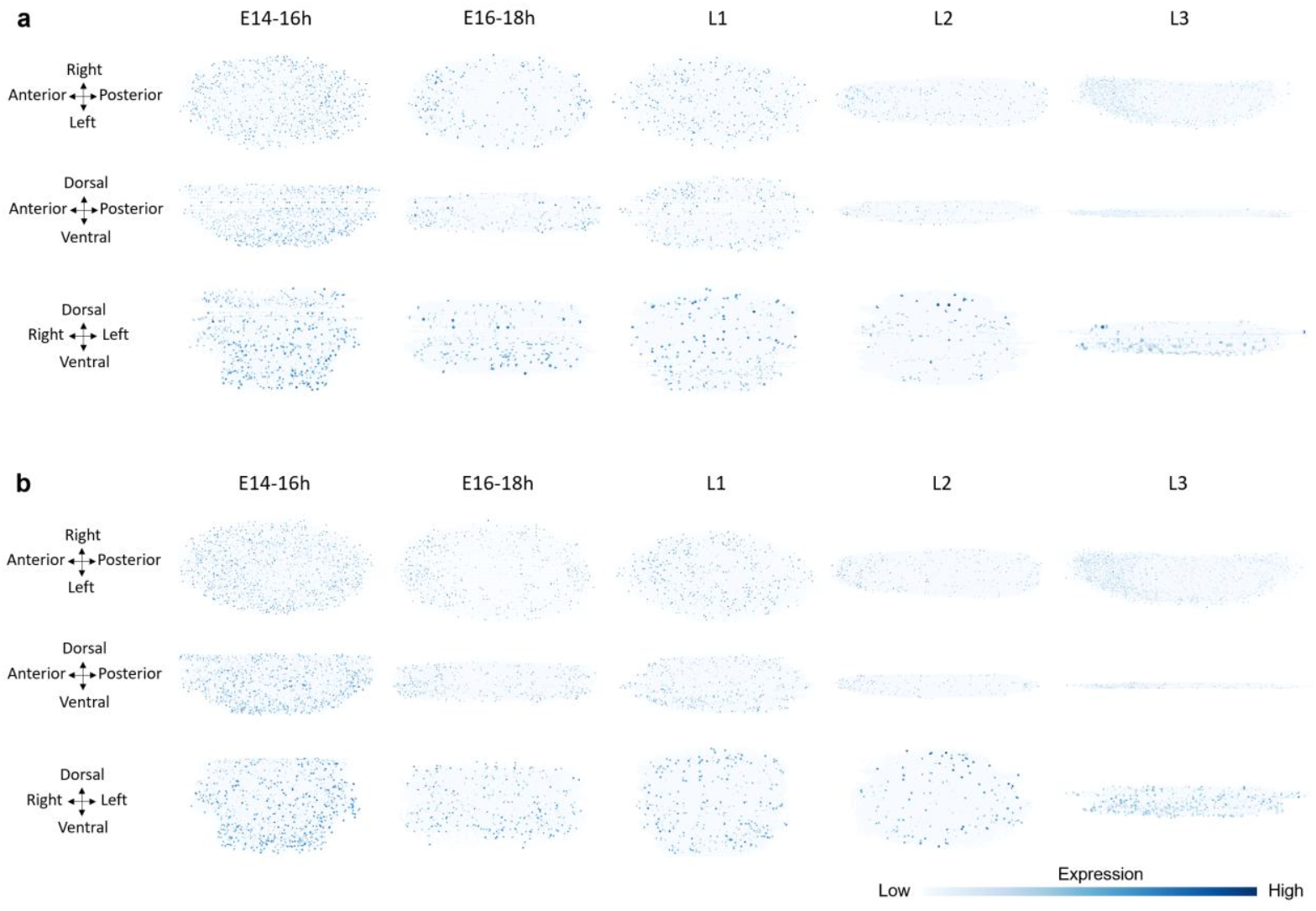
Additional Drosophila developmental comparisons. **a**, Raw low-resolution pseudo-3D stack of grh across developmental stages in different 3D views. **b**, SpatialZ 3D reconstruction of grh across developmental stages in different 3D views.

**Extended Data Figure 4:**
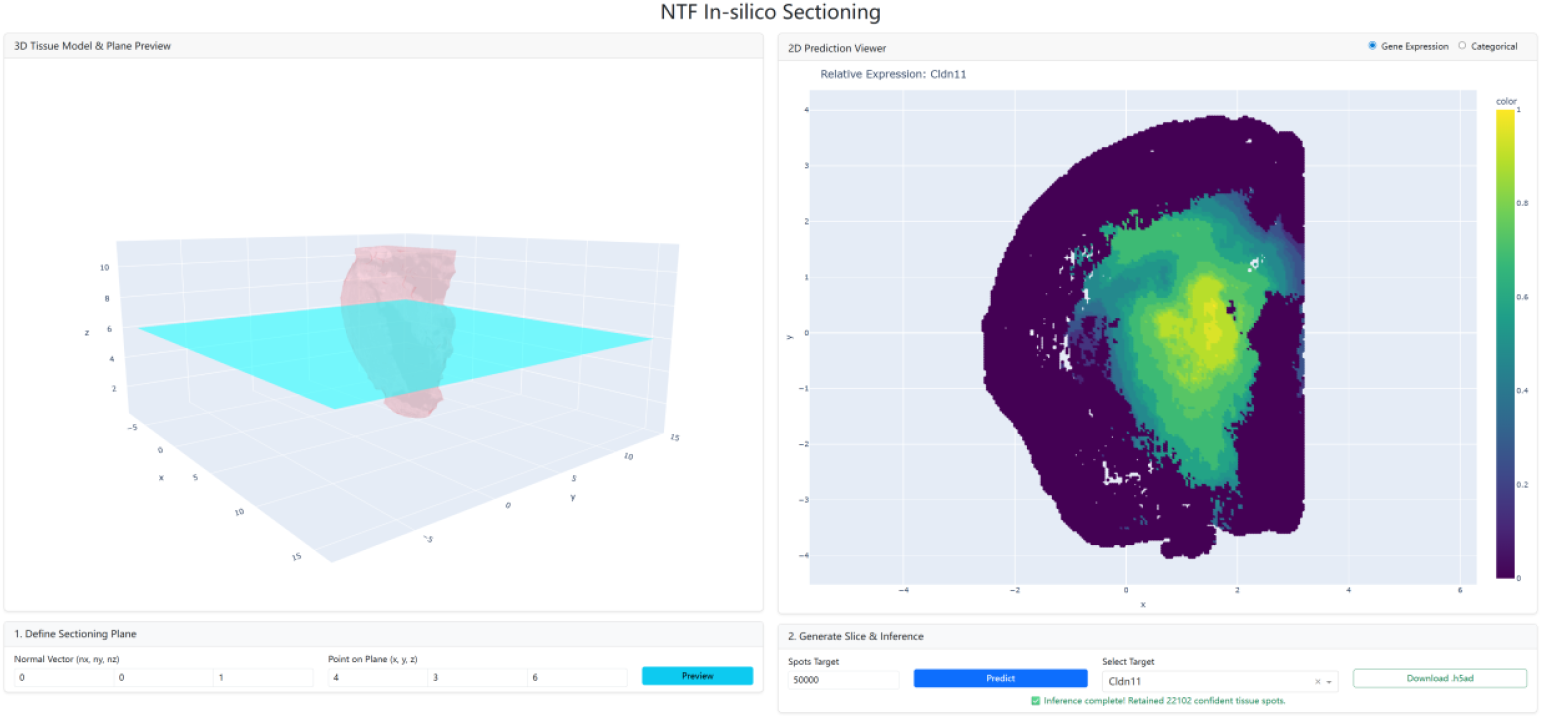
An example of NTF in silico sectioning app.

## REFERENCE

1 Crosetto, N., Bienko, M. & van Oudenaarden, A. Spatially resolved transcriptomics and beyond. Nat Rev Genet 16, 57–66 (2015). 10.1038/nrg3832

2 Moses, L. & Pachter, L. Museum of spatial transcriptomics. Nat Methods 19, 534–546 (2022). 10.1038/s41592-022-01409-2

3 Bressan, D., Battistoni, G. & Hannon, G. J. The dawn of spatial omics. Science 381, eabq4964 (2023). 10.1126/science.abq4964

4 Tian, L., Chen, F. & Macosko, E. Z. The expanding vistas of spatial transcriptomics. Nat Biotechnol 41, 773–782 (2023). 10.1038/s41587-022-01448-2

5 You, Y. et al. Systematic comparison of sequencing-based spatial transcriptomic methods. Nat Methods 21, 1743–1754 (2024). 10.1038/s41592-024-02325-3

6 Stahl, P. L. et al. Visualization and analysis of gene expression in tissue sections by spatial transcriptomics. Science 353, 78–82 (2016). 10.1126/science.aaf2403

7 Wang, X. et al. Three-dimensional intact-tissue sequencing of single-cell transcriptional states. Science 361 (2018). 10.1126/science.aat5691

8 Schott, M. et al. Open-ST: High-resolution spatial transcriptomics in 3D. Cell 187, 3953–3972 e3926 (2024). 10.1016/j.cell.2024.05.055

9 Sui, X. et al. Scalable spatial single-cell transcriptomics and translatomics in 3D thick tissue blocks. Nat Methods 22, 2574–2584 (2025). 10.1038/s41592-025-02867-0

10 Khan, M., Arslanturk, S. & Draghici, S. A comprehensive review of spatial transcriptomics data alignment and integration. Nucleic Acids Res 53 (2025). 10.1093/nar/gkaf536

11 Zeira, R., Land, M., Strzalkowski, A. & Raphael, B. J. Alignment and integration of spatial transcriptomics data. Nat Methods 19, 567–575 (2022). 10.1038/s41592-022-01459-6

12 Liu, X., Zeira, R. & Raphael, B. J. Partial alignment of multislice spatially resolved transcriptomics data. Genome Res 33, 1124–1132 (2023). 10.1101/gr.277670.123

13 Clifton, K. et al. STalign: Alignment of spatial transcriptomics data using diffeomorphic metric mapping. Nat Commun 14, 8123 (2023). 10.1038/s41467-023-43915-7

14 Jones, A., Townes, F. W., Li, D. & Engelhardt, B. E. Alignment of spatial genomics data using deep Gaussian processes. Nat Methods 20, 1379–1387 (2023). 10.1038/s41592-023-01972-2

15 Zhou, X., Dong, K. & Zhang, S. Integrating spatial transcriptomics data across different conditions, technologies and developmental stages. Nat Comput Sci 3, 894–906 (2023). 10.1038/s43588-023-00528-w

16 Lin, S. et al. Bridging the dimensional gap from planar spatial transcriptomics to 3D cell atlases. Nat Methods 23, 360–372 (2026). 10.1038/s41592-025-02969-9

17 Chen, Y. et al. From features to slice: parameter-cloud modeling of spatial transcriptomics for simulation and 3D interpolatory augmentation. bioRxiv (2025). 10.64898/2025.12.04.692251

18 Nagendran, M. et al. 1457 Visium HD enables spatially resolved, single-cell scale resolution mapping of FFPE human breast cancer tissue. Journal for ImmunoTherapy of Cancer 11, A1620–A1620 (2023). 10.1136/jitc-2023-SITC2023.1457

19 Janesick, A. et al. High resolution mapping of the tumor microenvironment using integrated single-cell, spatial and in situ analysis. Nat Commun 14, 8353 (2023). 10.1038/s41467-023-43458-x

20 Gong, Y., Yuan, X., Jiao, Q. & Yu, Z. Unveiling fine-scale spatial structures and amplifying gene expression signals in ultra-large ST slices with HERGAST. Nat Commun 16, 3977 (2025). 10.1038/s41467-025-59139-w

21 Mildenhall, B. et al. Nerf: Representing scenes as neural radiance fields for view synthesis. Communications of the ACM 65, 99–106 (2021).

22 Sitzmann, V., Martel, J., Bergman, A., Lindell, D. & Wetzstein, G. Implicit neural representations with periodic activation functions. Advances in neural information processing systems 33, 7462–7473 (2020).

23 Müller, T., Evans, A., Schied, C. & Keller, A. Instant neural graphics primitives with a multiresolution hash encoding. ACM transactions on graphics (TOG) 41, 1–15 (2022).

24 Wei, S. et al. Charting the spatial transcriptome of the human cerebral cortex at single-cell resolution. Nature Communications 16, 7702 (2025).

25 Wang, M. et al. High-resolution 3D spatiotemporal transcriptomic maps of developing Drosophila embryos and larvae. Dev Cell 57, 1271–1283 e1274 (2022). 10.1016/j.devcel.2022.04.006

26 Tomancak, P. et al. Global analysis of patterns of gene expression during Drosophila embryogenesis. Genome Biol 8, R145 (2007). 10.1186/gb-2007-8-7-r145

27 Mace, K. A., Pearson, J. C. & McGinnis, W. An epidermal barrier wound repair pathway in Drosophila is mediated by grainy head. Science 308, 381–385 (2005).

28 Almeida, M. S. & Bray, S. J. Regulation of post-embryonic neuroblasts by Drosophila Grainyhead. Mech Dev 122, 1282–1293 (2005). 10.1016/j.mod.2005.08.004

29 Kuett, L. et al. Three-dimensional imaging mass cytometry for highly multiplexed molecular and cellular mapping of tissues and the tumor microenvironment. Nat Cancer 3, 122–133 (2022). 10.1038/s43018-021-00301-w

30 Ribeiro, A. S. et al. P-cadherin functional role is dependent on E-cadherin cellular context: a proof of concept using the breast cancer model. J Pathol 229, 705–718 (2013). 10.1002/path.4143

31 Bologna-Molina, R., Mosqueda-Taylor, A., Molina-Frechero, N., Mori-Estevez, A. D. & Sánchez-Acuña, G. Comparison of the value of PCNA and Ki-67 as markers of cell proliferation in ameloblastic tumor. Medicina oral, patologia oral y cirugia bucal 18, e174 (2012).

32 Chen, A. et al. Spatiotemporal transcriptomic atlas of mouse organogenesis using DNA nanoball-patterned arrays. Cell 185, 1777–1792 e1721 (2022). 10.1016/j.cell.2022.04.003

33 Lein, E. S. et al. Genome-wide atlas of gene expression in the adult mouse brain. Nature 445, 168–176 (2007). 10.1038/nature05453

34 Caire, R. et al. A 3D morphogenetic blueprint for metastatic outgrowth in breast cancer. Cell (2026). 10.1016/j.cell.2026.03.009

