## Supplementary Table 3, Supplementary Figures 1-5 for "Ultra-efficient High Resolution 3D Reconstruction of Spatial Omics Data with Neural Transcriptomic Field"

### Supplementary Information

#### Supplementary Tables

**Table S3. 3D reconstruction performance (mean/sd) of different methods across different scenario in mouse visual cortex data**

| Methods | Spearman Correlation | RMSE | 3D SSIM |
|---|---|---|---|
| NTF (Scenario 1) | 0.48(0.02) | 1.00(0.26) | 0.86(0.06) |
| SpatialZ (Scenario 1) | 0.42(0.04) | 1.23(0.33) | 0.82(0.06) |
| FEAST (Scenario 1) | 0.32(0.05) | 1.76(0.21) | 0.69(0.04) |
| NTF (Scenario 2) | 0.44(0.04) | 0.96(0.27) | 0.94(0.04) |
| SpatialZ (Scenario 2) | 0.38(0.06) | 1.08(1.33) | 0.93(0.05) |
| FEAST (Scenario 2) | 0.23(0.03) | 1.85(0.22) | 0.82(0.04) |
| NTF (Scenario 3) | 0.52(0.04) | 1.01(0.24) | 0.89(0.07) |
| NTF (Scenario 4) | 0.40(0.05) | 0.99(0.25) | 0.97(0.02) |
| NTF (Scenario 5) | 0.38(0.02) | 1.01(0.23) | 0.92(0.04) |

### Supplementary Figures

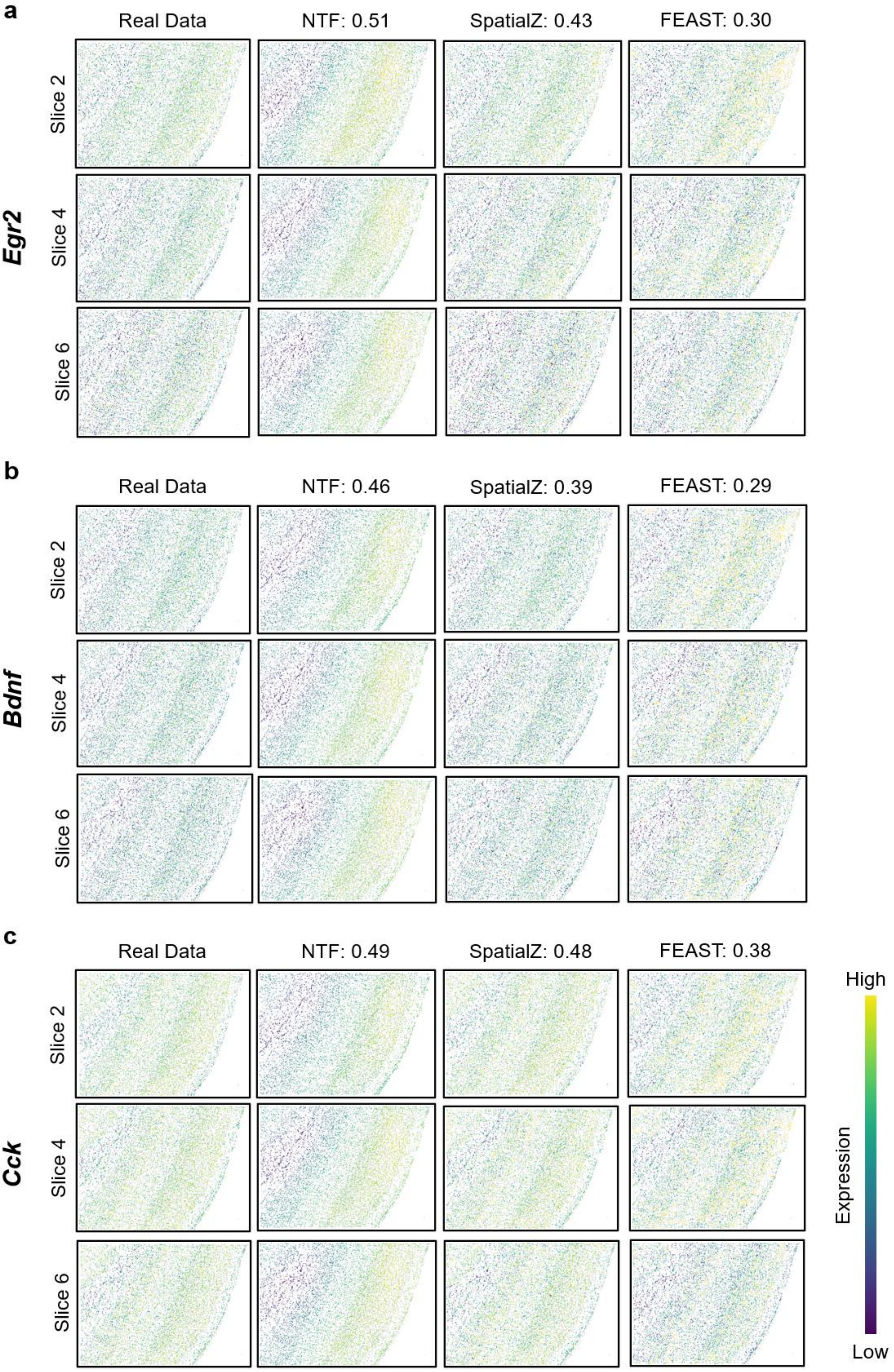

**SFig. 1 | Visualization of additional genes in intermediate-slice prediction setting on mouse visual cortex data.** In the first setting, we trained on slices 1, 3, 5 and 7 and reconstructed the held-out intermediate slices 2, 4 and 6. **a,** Ground truth and reconstruction results of gene *Egr2*. **b,** Ground truth and reconstruction results of gene *Bdnf*. **c,** Ground truth and reconstruction results of gene *Cck*.

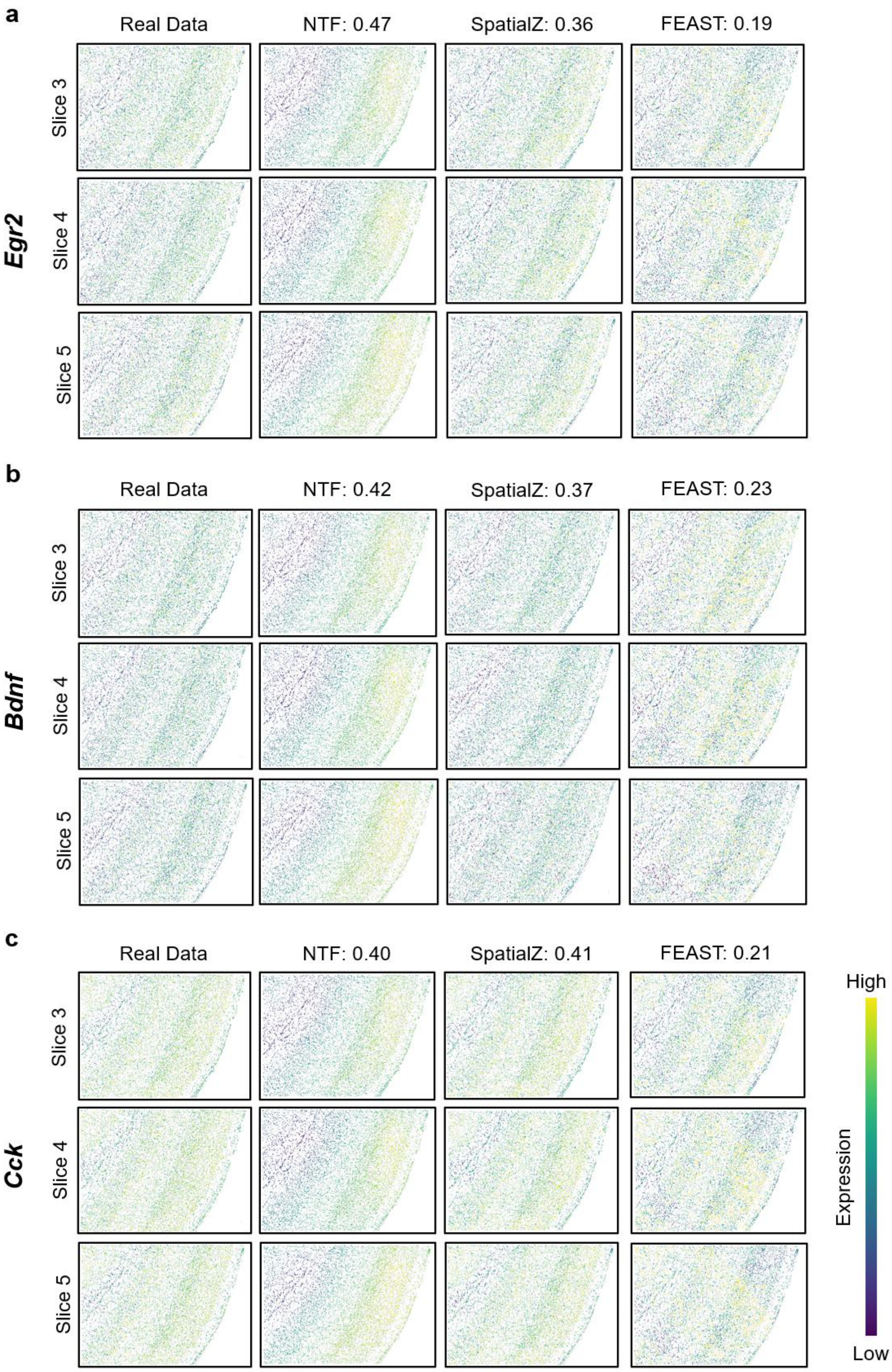


**SFig. 2 | Visualization of additional genes in large-block prediction setting on mouse visual cortex data.** Wider-gap held-out strategy in which slices 1, 2, 6 and 7 are used for training and slices 3, 4 and 5 are reconstructed. **a,** Ground truth and reconstruction results of gene *Egr2*. **b,**

Ground truth and reconstruction results of gene *Bdnf*. **c,** Ground truth and reconstruction results of gene *Cck*.

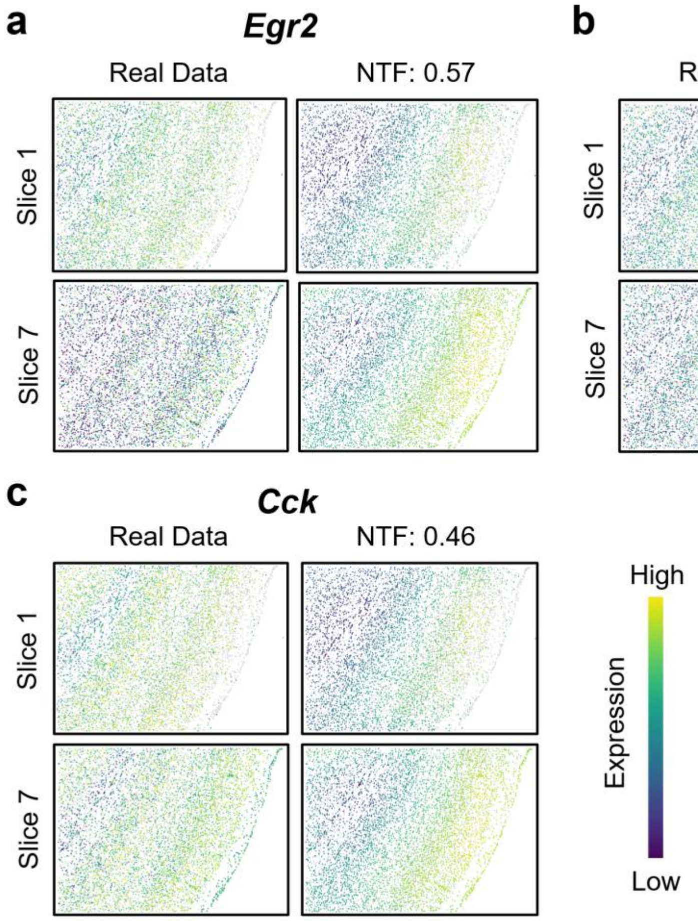


**SFig. 3 | Visualization of additional genes in two-side prediction setting on mouse visual cortex data.** This is extrapolative next-slice prediction setting in which slices 2-6 are used for training and slices 1 and 7 are predicted. **a,** Ground truth and reconstruction results of gene *Egr2*. **b,** Ground truth and reconstruction results of gene *Bdnf*. **c,** Ground truth and reconstruction results of gene *Cck*.

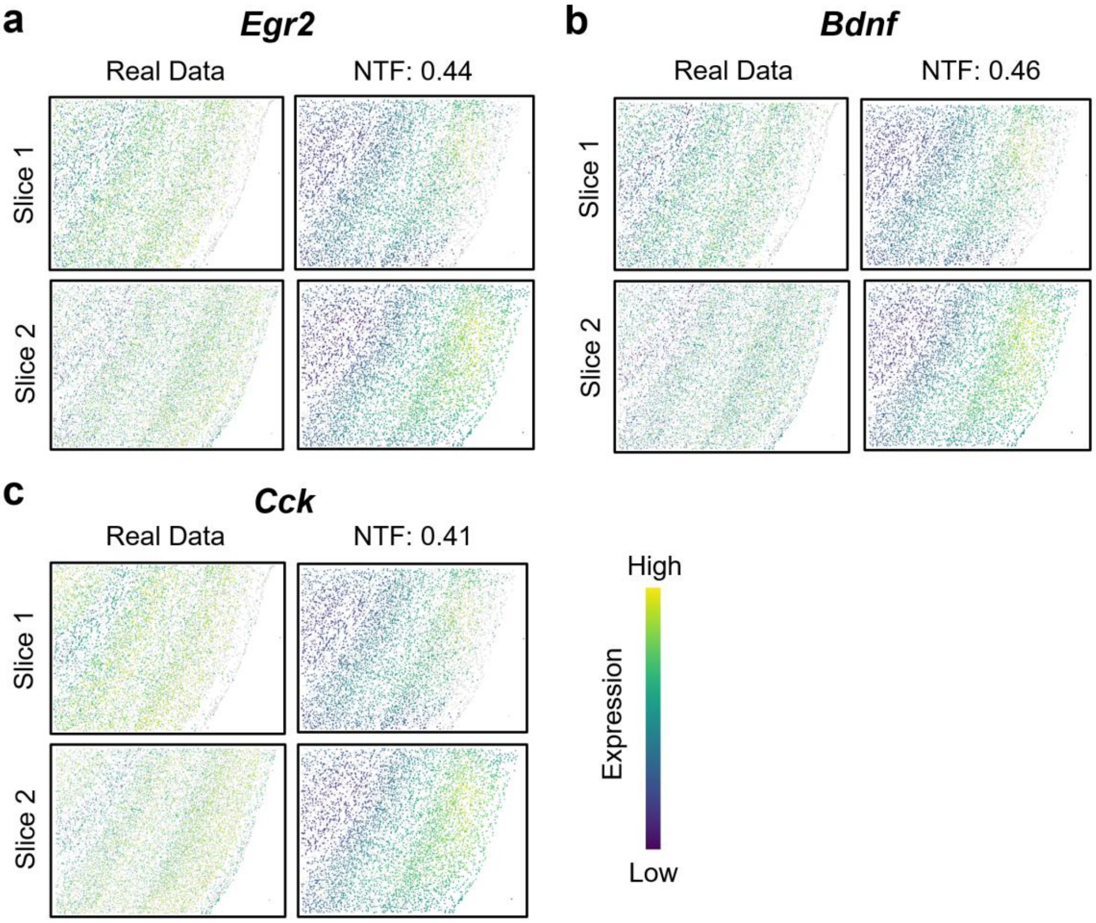


**SFig. 4 | Visualization of additional genes in head slice prediction setting on mouse visual cortex data.** This is backward extrapolation: NTF predicts the expression patterns of slices 1 and 2 using only slices 3–7 as training input. **a,** Ground truth and reconstruction results of gene *Egr2*. **b,** Ground truth and reconstruction results of gene *Bdnf*. **c,** Ground truth and reconstruction results of gene *Cck*.

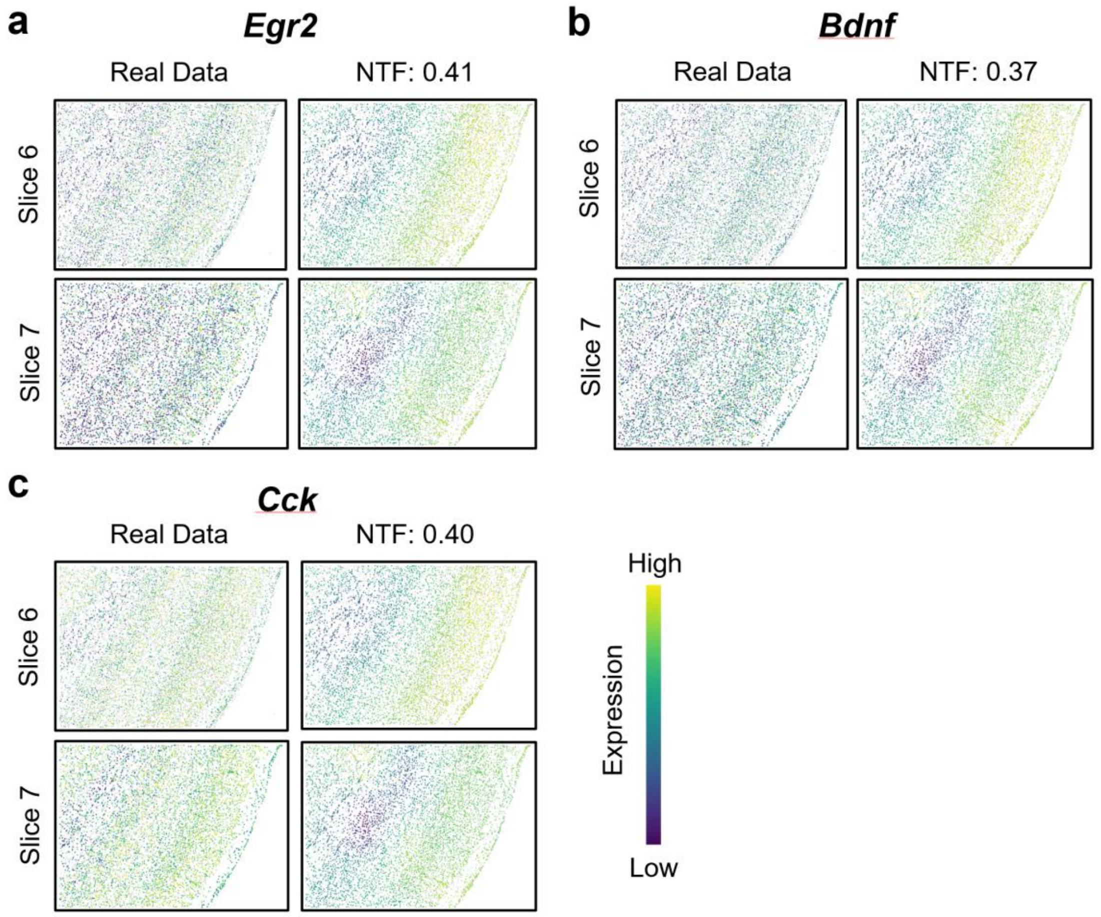


**SFig. 5 | Visualization of additional genes in tail slice prediction setting on mouse visual cortex data.** This is forward extrapolation: NTF predicts slices 6 and 7 using only slices 1–5 as training input. **a,** Ground truth and reconstruction results of gene *Egr2*. **b,** Ground truth and reconstruction results of gene *Bdnf*. **c,** Ground truth and reconstruction results of gene *Cck*.
